# Dynamic assembly of pentamer-based protein nanotubes

**DOI:** 10.1101/2024.08.15.608090

**Authors:** Lukasz Koziej, Farzad Fatehi, Marta Aleksejczuk, Matthew J. Byrne, Jonathan G. Heddle, Reidun Twarock, Yusuke Azuma

## Abstract

The molecular mechanisms and the geometrical theory underlying the polymorphic behavior of protein cages provide a basis for designing ones with the desired morphology and assembly properties. We show here that a circularly permuted cage-forming enzyme can controllably assemble into a variety of hollow spherical and cylindrical structures composed entirely of pentamers. A dramatic cage-to-tube transformation is facilitated by an untethered α-helix domain that prevents the 3-fold symmetry interaction and imparts a torsion between the building blocks. The unique double- and triple-stranded helical arrangements of subunits are mathematically optimal tiling patterns for this type of pentamer. These structural insights afford guidelines for the design of customized protein nanotubes for smart delivery and nanoreactor systems.

---

Engineering of biomolecular assemblies with precisely defined structure and functionality is the ultimate goal of bionanotechnology. While nucleic acids are the favored materials in the field (*1*), substantial efforts have been directed to the design of nanoarchitectures based on protein building blocks (*2*). Hollow spherical or cylindrical structures, called protein cages (*3-11*) or nanotubes (*12-15*), respectively, are particularly interesting in this context because of their prospective applications in delivery (*16-18*), catalysis (*19-21*), and nanomaterial construction (*22, 23*). Naturally occurring protein assemblies, such as viral capsids, provide design concepts as well as reengineering platforms for customized nanodevice development (*24, 25*).

Many viral coat proteins can assemble into particles with a range of sizes and shapes when reconstituted *in vitro* (*26, 27*). Geometrical patterns defined by quasi-equivalence (Casper-Klug) theory (*28*) explain such polymorphic behavior. The variable number of hexamers filling in the gap between pentameric subunits at the vertices results in cage expansions and irregular forms (*29, 30*). Such a dynamic, polymorphic feature is useful for customizing assemblies (*27, 31*), as demonstrated with the cowpea chlorotic mottle virus (CCMV) coat protein, which assembles around DNA origami structures and protects them from degradation (*32*).

Unique polymorphic behavior has been observed for engineered variants of *Aquifex aeolicus* lumazine synthase (*33, 34*). While the wild-type protein, AaLS-wt, self-assembles into a ∼16-nm dodecahedral structure composed of 60 identical monomers (Fig. 1A) (*35*), the negatively supercharged variants AaLS-neg and AaLS-13 (*36, 37*) adopt expanded 360- and 720-mer assemblies, respectively, constructed entirely from pentameric capsomers (*38*). Moreover, a circularly permuted variant of AaLS, cpAaLS(119) (Fig. 1B), forms not only ∼24-nm and ∼28-nm expanded spherical cages, but also straight ∼24-nm-wide tubes of variable length (*39*).

**Fig. 1.**
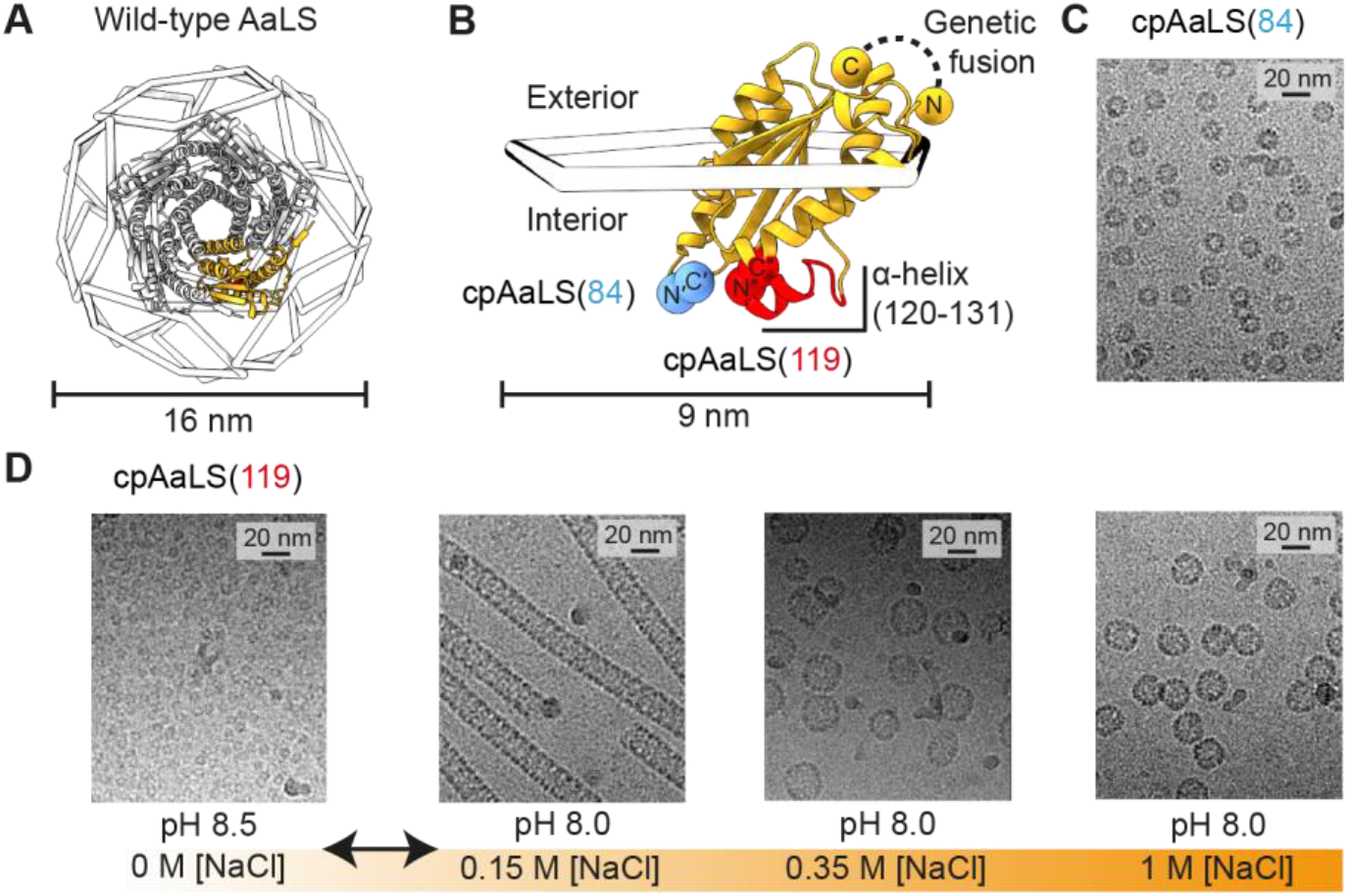
Assembly control of circularly permuted AaLS. (**A**) Structure of the dodecahedral wild-type AaLS cage (PDB ID 1HQK), shown as 12 wire pentagons with a ribbon diagram of a representative pentamer (grey) and protomer (orange). (**B**) Design of the circularly permuted variants, cpAaLS(84) and cpAaLS(119). The peptide linker connecting the native N- and C-termini (GTGGSGSS) is shown with a black dashed line. The new termini, C′(84) and N′(85) (blue) or C″(119) and N″(120) (red), are indicated by spheres. The α-helix(120-131), untethered by circular permutation for cpAaLS(119), is highlighted in red. (**C**,**D**) Cryo-EM micrographs of the cpAaLS(84) cage (C) and the NaCl- and pH-dependent cpAaLS(119) assemblies (D).

Although such characteristics potentially present novel design principles (*31*), the geometric blueprints and the molecular mechanisms underlying this polymorphic behavior have remained unknown. Using cryo-EM and mathematical modeling, we here elucidate how pentameric building blocks can controllably and dynamically assemble into specific spherical and tubular structures in preference to the other possible particle morphologies.

## Salt- and pH-dependent assembly of cpAaLS(119)

We previously designed two circularly permuted variants of AaLS and confirmed that their morphologies depend on the positions of the newly generated N- and C-termini (*39-41*). The native terminal amino acids were connected via an octapeptide linker using genetic fusion and new sequence termini were introduced either between residues 84 and 85 or 119 and 120, yielding cpAaLS(84) or cpAaLS(119), respectively (Fig. 1B). While cpAaLS(119) exhibits polymorphic behavior as discussed above, the cpAaLS(84) protein forms a ∼16-nm diameter homogeneous cage structure, similar to AaLS-wt, and serves as a control for the experiments described below (Fig. 1C).

While characterizing the cpAaLS(119) variant, we unexpectedly found that the cage-like structures disassemble into fragments at low ionic strength and alkaline conditions. The protein was heterologously produced in *Escherichia coli* and isolated using ion exchange chromatography. Subsequent buffer exchange to 5 mM Tris-HCl buffer at pH 8.5 resulted in almost completely disassembled fragments, confirmed by size-exclusion chromatography coupled with right/low-angle light scattering detectors (SEC-RALS/LALS) (fig. S1).

Isolation of the cpAaLS(119) capsomers enabled systematic investigation of the reassembly process. Cage fragments were subjected to a rapid buffer exchange to 50 mM Tris-HCl buffer at pH 8.0 containing varying concentrations of NaCl (fig. S2A), and the resulting assemblies were analyzed by SEC and cryo-electron microscopy (cryo-EM) (Fig. 1D, and fig. S2, B to E). While remaining as unassembled fragments in the absence of salt (fig. S2B), adding 0.15 M NaCl facilitated cpAaLS(119) tubular formation with ∼92% yield (fig. S2C). Further increasing NaCl concentration to 0.35 M yielded a mixture of the tubes and ∼28-nm spherical cages ∼44% and ∼46%, respectively (fig. S2D). In 1 M NaCl, the tubes remained a 2% minority, while the ∼28-nm cage constituted ∼30% and the smaller ∼24-nm cages dominated at ∼59% (fig. S2E). These results demonstrated that cpAaLS(119) assembly can be controlled by adjusting the ionic strength of the solution. As NaCl concentration increases, unassembled fragments are preferentially transformed into 24-nm wide tubes, as well as 28-nm-, and 24-nm spheres.

Small changes in buffer pH can modulate the salt-dependent assembly of cpAaLS(119). Lowering pH tends to favor the assemblies that appear at high ionic strength and *vice versa*. For example, with 1 M NaCl at pH 8.5, 8.0, and 7.5, the proportion of 24-nm spherical cages was 9%, 59%, and 74%, respectively (fig. S2E). Furthermore, salt/pH-dependent cpAaLS(119) cage assembly is fully reversible. When the tube and spherical cages were subjected to buffer exchange into 50 mM Tris-HCl buffer at pH 8.5, the protein was completely converted into disassembled cage fragments (fig. S3).

To test if the cpAaLS variants retained the extreme thermal stability from the parent AaLS-wt, which has a melting temperature (*T*_m_) greater than 120°C (*35, 42*), we performed a thermal shift assay based on tryptophan fluorescence, coupled with dynamic light scattering, at different pH and NaCl concentrations (fig. S4A). Irrespective of the tested buffer conditions, cpAaLS(84) did not exhibit substantial fluorescent changes up to 110°C, indicating little protein unfolding (fig. S4B). Temperature-dependent protein denaturation was observed for cpAaLS(119) with a *T*_m_ of 88-103°C, where higher ionic strength and lower pH tend to increase the thermal stability. This variant also showed a decrease in size, probably due to partial fragmentation, at 70-80°C under conditions that promote tube formation (fig. S4C, pH 7.5 and 0.15 M NaCl). Topological and/or morphological alteration likely leads to a relatively lower heat tolerance of cpAaLS(119) than that of AaLS-wt.

## Geometric blueprints of the cpAaLS assemblies

Control over cpAaLS(119) assembly by salt and pH allowed us to determine the structures of individual morphologies using cryo-EM single particle and helical reconstruction (Fig. 2, and figs. S5 and S6). As previously hypothesized (*39*), all the structures were found to consist exclusively of the pentameric building blocks. The cpAaLS(84) assembly resembles the AaLS-wt cage, where each pentamer interacts with five neighboring subunits in a dodecahedral arrangement (fig. S5A). In contrast, cpAaLS(119) assemblies are non-quasi equivalent, where at least 1 interface of the pentameric subunits remains uncontacted. The 24-nm and 28-nm spherical cages are composed of 24- and 36-pentamers, respectively, and both have tetrahedral symmetry (Fig. 2, C and D), resembling those formed by the previously engineered AaLS variants, NC-1 and AaLS-neg (*38, 43*). In the tubular structure, the building blocks are arranged as a triple-stranded helix (Fig. 2B), which is unique and has never been seen for any natural or engineered proteins.

**Fig. 2.**
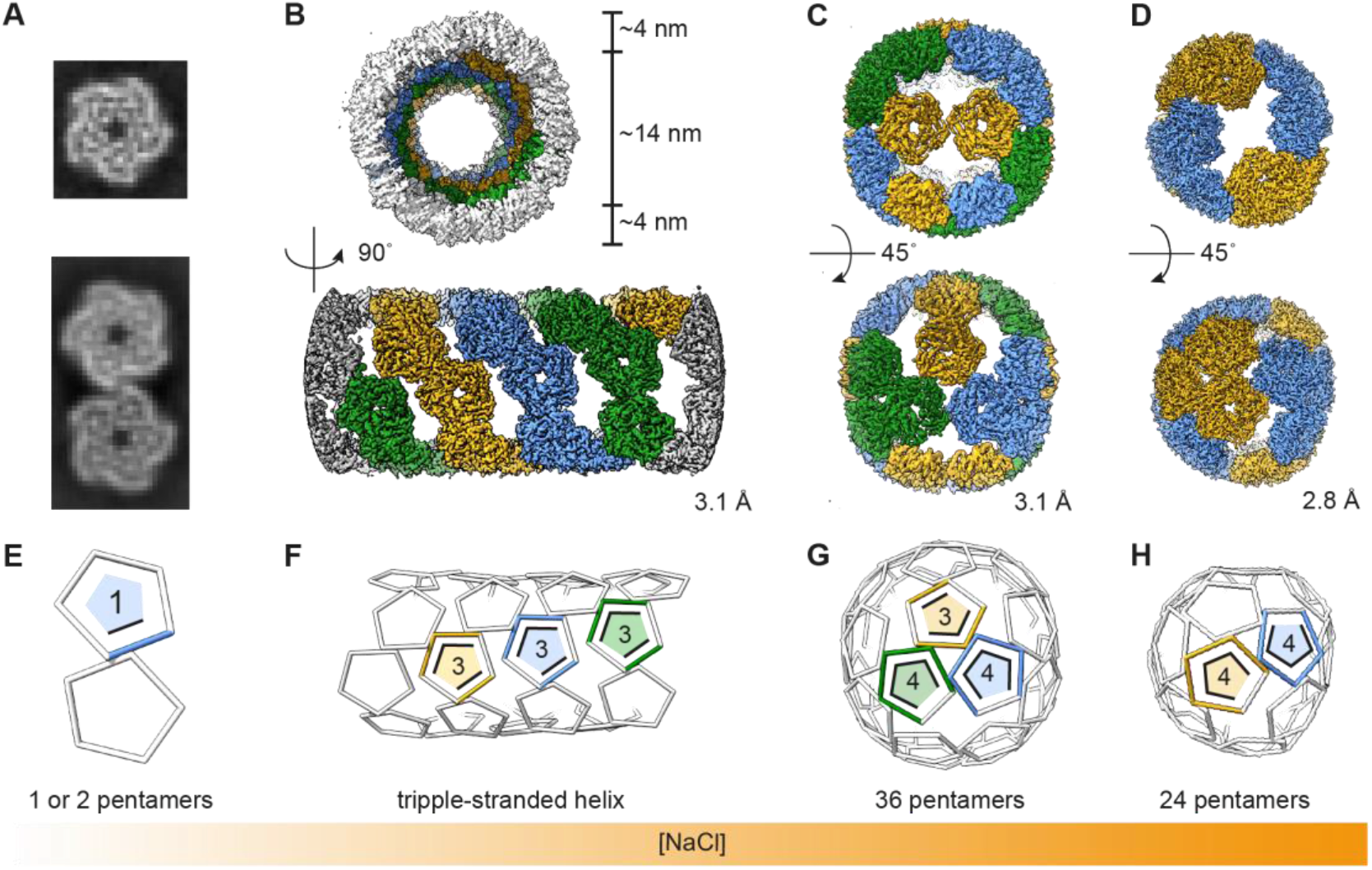
Cryo-EM structures of the cpAaLS(119) assemblies. (**A-D**) 2D classes (A) and 3D maps (B-D) of the cpAaLS assemblies, where colors (blue, orange, or green) indicate individual threads in the helical structure (B) or symmetry-related pentameric subunits in the spherical cages (C,D). The resolution of the final 3D reconstructions (GS-FSC at 0.143 cut-off) is provided at the right corner of each map. (**E**-**H**) The corresponding wire representation of the cpAaLS assemblies with the number of contacts per each asymmetric pentamer. Images are not to scale.

The transformation of cpAaLS assemblies is accompanied by changes in the number of connections between the constituent pentamers. While the number of pentamer-pentamer contacts remains 0 or 1 without NaCl at pH 8.5, judged by the 2D-averaged cryo-EM images (Fig. 2, A and E), the mean contact number per pentamer increases to 3, 3.7, and 4 in the tubular, 28-nm-, and 24-nm spherical cages, respectively (Fig. 2, F to H). This trend suggests that increasing salt and/or lowering pH stabilizes inter-pentamer interactions.

AaLS is known to have a higher number of ionic interactions and hydrogen bonds at the pentamer-pentamer interface compared to an analogous lumazine synthase derived from *Bacillus subtilis* (*35*). These charge-driven interactions have been hypothesized to contribute to its extreme thermal stability. However, the salt-dependent increase in the number of contacts observed for cpAaLS(119) structures suggests that hydrophobic interactions are the major driving force for their assembly. Meanwhile, the pH-dependency can be explained by the net negative charge of AaLS protein, which has a theoretical isoelectric point of 5.8. An increase in pH potentially endows the pentamers with increased negative surface charge, weakening inter-pentameric interactions due to charge repulsion.

## Molecular mechanisms underlying the polymorphic behavior

The atomic model of the cpAaLS(84) cage revealed that all the constituent pentamers interact with each other in the same manner as in the AaLS-wt assembly (Fig 3A). The 2-fold symmetrical interface consists of a hydrophobic patch (L8, L141, and W137) surrounded by a hydrogen bond (H41) and ionic interactions (e.g. R40 and E5) (fig. S7, A and B) while an α-helix region (120-131) forms 3-fold symmetric hydrophobic clusters (I121 and I125) in the cage interior (Fig 3, B and C).

**Fig. 3.**
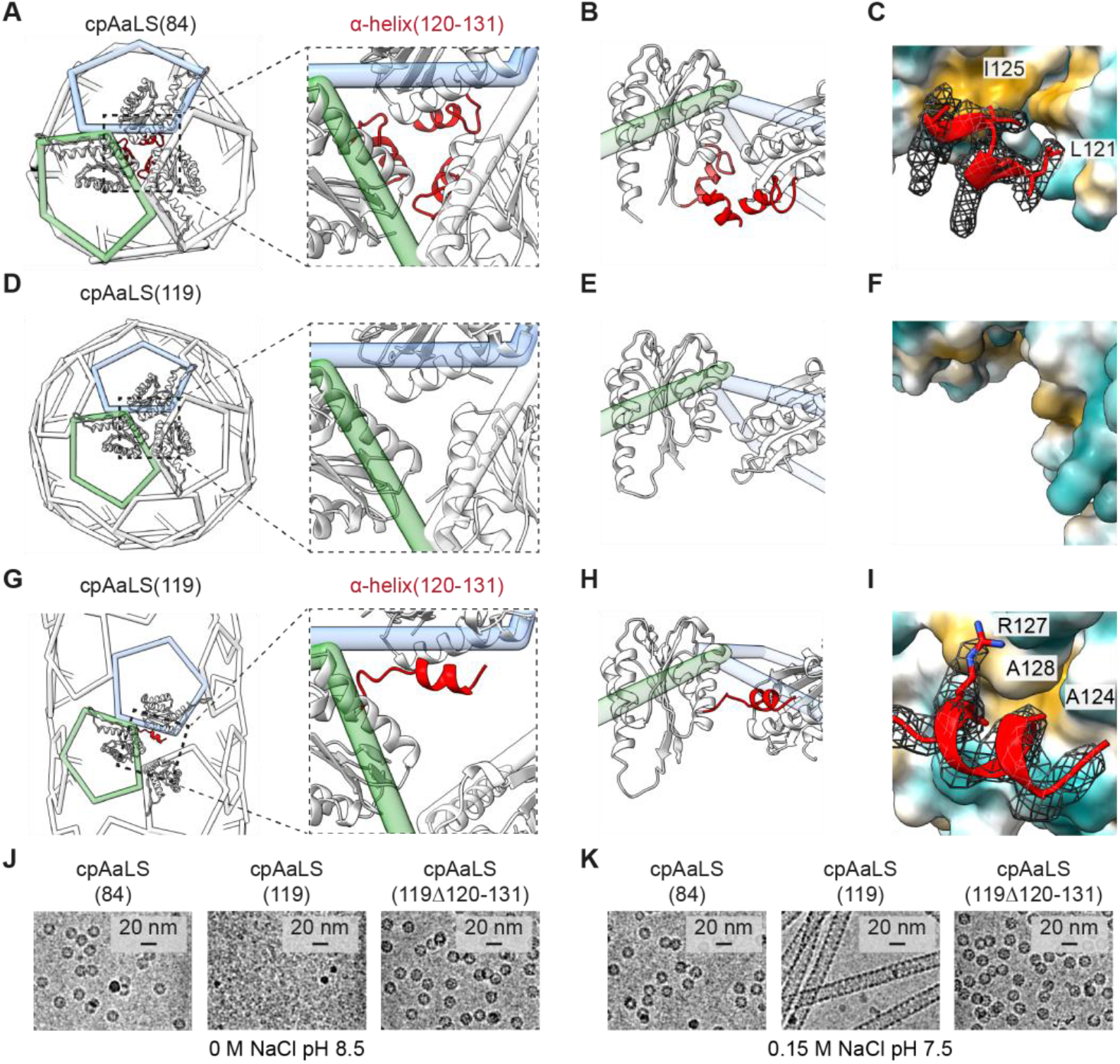
‘Untethered’ α-helix(120-131) facilitating the dynamic assembly of cpAaLS(119). (**A**) Wire diagram of the cpAaLS (84) assembly with an enlarged view of the 3-fold symmetry region. Three interacting monomers are shown as a ribbon with α-helix(120-131) highlighted in red. (**B**) Rotated side view of a pentamer pair (green and blue wire). The α-helix(120-131) domain from another monomer at the front is also shown to present the interaction at the 3-fold symmetry region in the cpAaLS(84) cage lumen. (**C**) Atomic interaction mode of the α-helix(120-131) domain in the cpAaLS(84) assembly. A unit is shown as a ribbon with amino acid side chains at the interface with the corresponding cryo-EM density map (mesh), and the interacting partners as hydrophilic (cyan) and hydrophobic (light brown) surfaces. (**D-I**) The corresponding representations for the cpAaLS(119) 24-pentameric spherical cage (D-F) and the tubular assembly (G-I), where the α-helix(120-131) is structurally disordered and was not modeled (D-F), or flipped to interact with an alternative surface (G-I), respectively. Panel (F) shows the same region as (C) to present the lack of cryoEM density corresponding to the α-helix(120-131) region. (**J**,**K**) Cryo-EM images of the cpAaLS(119) variant lacking α-helix(120-131), cpAaLS(119Δ120-131), compared to those of cpAaLS(84) and cpAaLS(119).

The bonding network observed at the 2-fold symmetry interface of AaLS-wt is approximately preserved for all the interacting pentamers in the cpAaLS(119) assemblies (fig. S7, C to F). There is no swapping in the amino acid interaction partners in this region upon circular permutation and morphology change. In marked contrast, the (pseudo) 3-fold symmetrical interaction interface of cpAaLS(119) assemblies lacks cryoEM density corresponding to the N-terminal α-helix(120-131) (Fig. 3, D to F). This finding suggests that the α-helix is unable to form the native-like hydrophobic cluster, likely due to the disconnection of residues 119 and 120 for circular permutation (Fig. 1B).

In cpAaLS(119) tubes, the ‘untethered’ α-helix(120-131) binds to the adjacent pentamer surface, which is a solvent-exposed region in wild-type-like assemblies (Fig. 3, G to I). Because of this non-native interaction, the α-helix(120-131) appears to block 3-fold symmetrical pentamer-pentamer interactions, which explains the uncontacted interfaces observed in the cpAaLS(119) assemblies (Fig. 2, E to H). Indeed, the deletion variant lacking the α-helix(120-131) domain, cpAaLS(119Δ120-131), does not exhibit polymorphic behavior, but assembles into only wild-type-like ∼16-nm spherical cages (Fig. 3, J and K). These results prove that the untethered α-helix(120-131) domain is essential for the non-native, expanded cage formation of cpAaLS(119).

In thermal shift assays, the cpAaLS(119Δ120-131) cage showed a similar behavior to cpAaLS(84): having no denaturation up to 110 °C (fig. S4). These results suggest that the reduced thermal stability of cpAaLS(119) is mainly due to the morphology, in which constituent pentamers lack one or two interactions with their neighbors, rather than the loss of the hydrophobic core in the region of the 3-fold symmetry axis.

The binding of the untethered α-helix(120-131) to neighboring pentamers appears to be weak and occasional. This is suggested by an atomic-level interaction mode in which an arginine (R127) and two alanine (A128 and A124) residues from the helical domain contact a serine in the linker connecting the native termini and a hydrophobic cleft formed between intra-pentameric monomers, respectively (Fig. 3I). Furthermore, substantial cryo-EM density corresponding to the α-helix(120-131) was found only for 2 protomers in each pentamer constituting the tubular assembly and invisible in the spherical assemblies. The dynamic nature of the binding, which partially blocks pentamer-pentamer interactions, likely reflects multiple assembly states of cpAaLS(119).

## Capsomer interaction angles facilitating the tubular formation

The unique triple helix is not the only tubular structure formed by cpAaLS(119). In the cryo-EM micrographs, short and bent caterpillar-like objects, referred to as “twisted tubes” were also observed albeit at a very low frequency (Fig. 4A, left). Due to the limited number of particles, helical reconstruction was not initially possible. However, introduction of C37S and A85C mutations into cpAaLS(119) to give cpAaLS(119, C37S, A85C) and a modified assembly protocol (fig. S8) surreptitiously resulted in enrichment of the twisted tubular structure and allowed subsequent cryo-EM analysis (Fig. 4A, right). Since the twisted tubular assemblies were heterogeneous, we divided particle images into 3 classes and determined the structures individually (fig. S9).

**Fig. 4.**
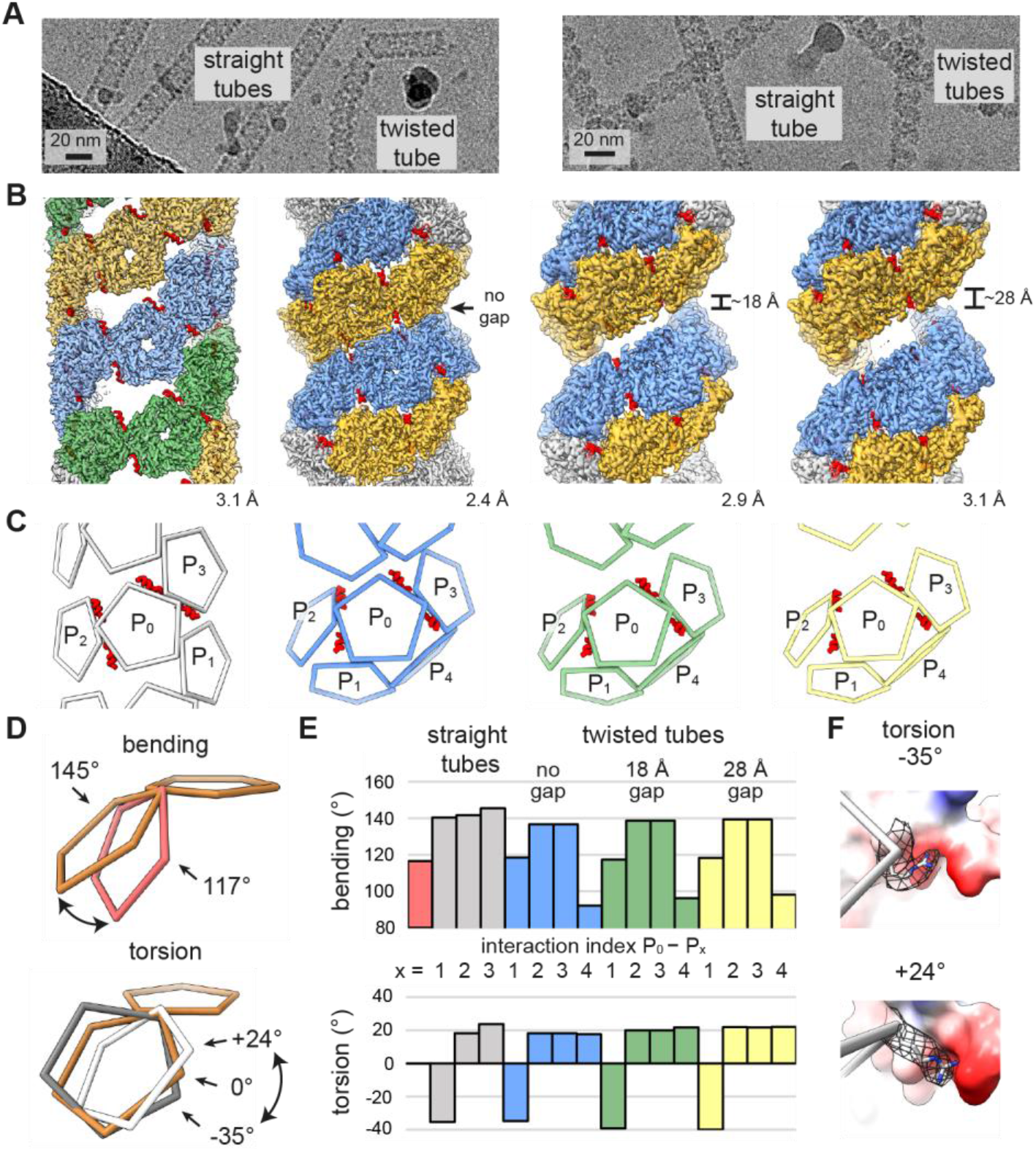
Capsomer interaction angles in the cpAaLS(119) tubes. (**A**) Cryo-EM micrographs of cpAaLS(119) (left) and cpAaLS(119, C37S, A85C) (right). (**B**) Cryo-EM maps of the straight tube composed of 3 evenly spaced helical strips (green, orange, and blue) and the twisted tubes featuring a variable gap (0, ∼18, or ∼28 Å) between dual strips (blue and orange). The resolution of the final 3D reconstructions (GS-FSC at 0.143 cut-off) is shown at the right corner of each map. The maps are not to scale. (**C**) The corresponding wire representations showing interactions of a pentamer (P_0_) with neighbors (P_1-4_). The α-helix(120-131) domains that accompany the intrathread interactions (P_0_-P_2/3_), otherwise invisible, are highlighted as red ribbons. (**D, E**) Conceptual representation (D) and the measured values (E) of the bending (top) and the torsion angles (bottom) between two pentamers. The wire diagrams show a pentamer-pentamer interaction in the cpAaLS(84) cage (red, top) and the cpAaLS(119) straight tubes (white and grey, bottom) with a reference having 0° torsion and 145° bending angles (brown). The interactions in the bar graph are colored and indexed as in (D), with those of the cpAaLS(84) cage (red bar) shown for comparison. (**F**) Two edges between interacting pentamers with external torsion angles (−35º top, 24º bottom) in the cpAaLS(119) straight tube. The arginine side chain at the right vertex (R40, shown as grey sticks with mesh for the corresponding cryo-EM map) is flipped to maintain its interaction with a negatively charged patch (red surface) in the opposite pentamer.

All three distinct twisted tube structures are composed of two compacted helical threads with 0-, 18-, and 28-Å gaps between them (Fig. 4B). This spring-like arrangement rationalizes the bending tendency. We also realized that one of the four pentamer interactions in the twisted tube has an acute bending angle (Fig. 4C, P_0_-P_4_ interaction). A possible disulfide bridge between two of the introduced cysteine residues at position 85 seems to support the unusual interaction mode, enhancing the twisted tubular formation (fig. S10).

As highlighted by the twisted tubes, the capsomer’s interaction angles and the entire morphology are interlinked. While each pentamer-pentamer interface in the regular AaLS-wt assembly is tightly fixed in a single pattern by the 2-fold and two 3-fold symmetry interactions, removal or modulation of these contacts by reengineering likely leads to a flexible connection and expanded cage-like structures. To analyze the relative positioning of the pentamers in the cpAaLS assemblies systematically, we built a script that calculates the bending and torsion angles of two interacting polygon-shaped multimers (Fig. 4D). We named the computational angle generator “AngelaR”.

Analysis of the cpAaLS spherical assemblies using AngelaR showed that an increase in the bending angles between interacting pentamers results in larger-sized structures, as reported previously (*38*) (fig. S11). In the twisted tubes, the increased gap between the compacted helical threads is accompanied by decreased bending and an increased torsion angle, a tendency agreeing with computational simulation using a pentagon-based helical model (fig. S12). In both the straight and twisted tubes, the pentamer-pentamer interaction adopts a wide range of torsion angles: -35º, 18º, and 24º for the straight tube, for example. The prominent negative torsion angles enable interthread dockings in these helical structures (Fig. 4, C and E, P_0_-P_1_ interaction).

AaLS proteins accept variable interaction angles using the flexible nature of the amino acid side chain. At the 2-fold symmetry interfaces of the cpAaLS(119) straight tube, for instance, an arginine residue (R40) is flipped to retain the interaction with the same negatively charged surface of the neighboring pentamer (Fig. 4F), being accompanied by the interaction torsion-angle change.

When focusing on the individual thread of the tubular assemblies, the constituent pentamers are connected through consistently positive torsion angles (Fig. 4, C and E, P_0_-P_2/3_ interactions). This is required for helical arrangements of this type of pentameric building blocks, and the torsion angle defines the helical rise. If the torsion angle is zero degrees, as in the case of AaLS-wt assembly, the pentamers can form only a ring-shaped arrangement with no helical rise (Fig 4E, red bar). Notably, these intrathread capsomer interactions involve a pair of the untethered α-helices(120-131) that bind with a neighbor on the flipped position (Fig. 4, B and C, P_0_-P_2/3_ interactions), probably supporting their torsion angles in a certain range.

The α-helix (120-131) domain is structurally disordered and unseen in cryo-EM analysis of the cpAaLS(119) spherical cages (Fig. 3, D to F). Tubular assemblies have never been observed for other AaLS variants in which the α-helix(120-131) is tethered in the native position (*38, 43-45*) or removed (Fig 3, J and K). Considering these observations together, we conclude that the untethered α-helix(120-131), which blocks the 3-fold symmetry interaction (Fig. 3G) while imparting a consistent torsion angle to the subunits (Fig. 4, D and F, P_0_-P_2/3_ interactions), is the essential element for the transformation of the AaLS assembly from spherical to tubular structures.

## Mathematical rationale for the pentagon-based tubular structures

The spectrum of protein assemblies formed by a specific building block can be classified via tiling theory (*29, 30, 46-49*). To gain geometrical insight into the cpAaLS(119) tubular assemblies, we simulated all the possible helical arrangements formed by pentagons on a planar lattice (Fig. 5A and fig. S13). The model consists of a pair of pentamers (red) arranged with vectors pointing along 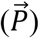 and across 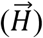 helical threads, where the geometrically possible tubular structures are characterized by the number of distinct strips (helicity number *n*_*h*_) and the translation steps along 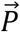 (periodicity number *n*_*p*_). Some structures defined by these parameters, e.g. (*n*_*h*_, *n*_*p*_) = (4, 4), require flipped pentagons, which can not be realized by asymmetric interaction surfaces of proteins (fig. S14). The cpAaLS(119) straight tubes adopt the tiling pattern with *n*_*h*_ = 3 and *n*_*p*_ = 4, one of the smallest structures among all the geometrically and biologically possible options (Fig. 5, A and B).

**Fig. 5.**
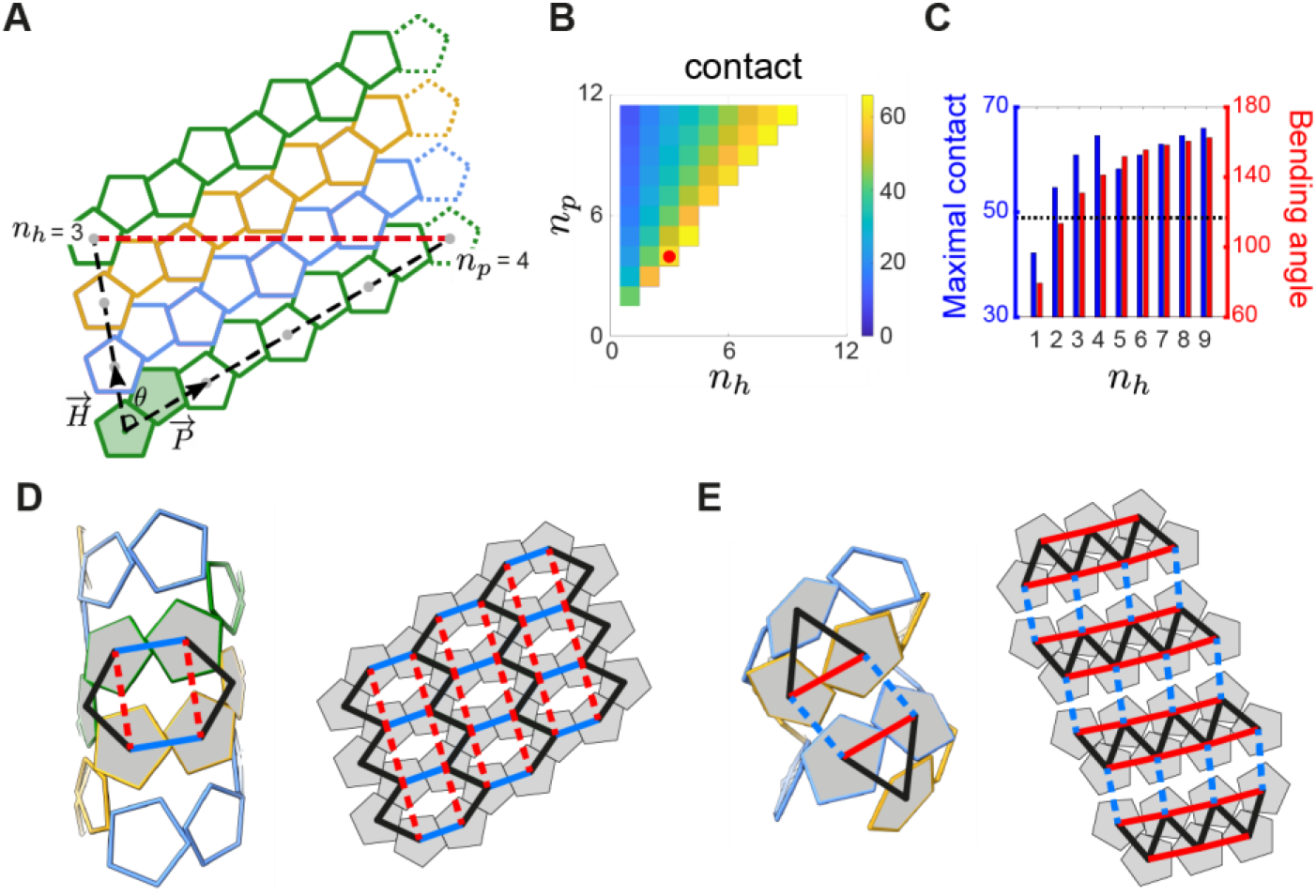
Geometric rationale for the cpAaLS(119) tubular assembly. (**A**) A tiling representation of the cpAaLS(119) straight tube composed of three pentamer strips (green, blue, and orange). The lattice model is formed from periodic repeats of pentamer pairs (shaded in green) with vectors pointing along 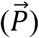 and across 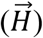 helical threads at an angle (*θ*). The tubular architecture is defined by the translation steps, helicity (*n*_*h*_) and periodicity (*n*_*p*_) numbers, for two pentamers distanced by a single helical turn (grey dots connected by a black dashed line). The pentamers connected with the red dashed line are identical in the 3D tubular structure. (**B**) Heatmap presenting the percentage contact area between pentagonal edges in the tubular model for different (*n*_*h*_, *n*_*p*_) combinations. The experimentally observed (*n*_*h*_, *n*_*p*_) = (3,4) is indicated by a red dot. The structures in the blank area are geometrically or biologically forbidden. (**C**) The maximally possible contact area (blue) and the bending angle (red) for given *n*_*h*_ over all the possible choices of *n*_*p*_. The bending angle for the wild-type-like cpAaLS(84) cage is shown as a dotted line. (**D-E**) The interaction network rewiring between pentamers (shaded in grey) in the straight (D) and twisted tube (E), superimposed onto the 3D models (left) and the 2D tiling (right). Solid and dashed lines indicate contact and non-contact between pentamers, respectively. In the network transformation from the straight to the twisted tube, the blue contacts were lost, while the red ones were gained.

The parameters (*n*_*h*_, *n*_*p*_) = (3, 4) appear to provide the optimal tiling pattern for the capsomer interaction of the AaLS protein. We next analyzed the percentage contact length between adjacent pentagons for different (*n*_*h*_, *n*_*p*_) combinations, finding that the maximal contact for each *n*_*h*_ is consistently obtained with the smallest possible *n*_*p*_ (Fig. 5B). Comparison of the maximal contact for different *n*_*h*_ identifies (*n*_*h*_, *n*_*p*_) = (4, 5) as the local maximum and (*n*_*h*_, *n*_*p*_) = (3, 4) as the second-best option among the smaller *n*_*h*_values (Fig. 5C, blue bars). We further benchmarked the bending angles between adjacent pentagons in the different geometric options (Fig. 5C, red bars), indicating that the (*n*_*h*_, *n*_*p*_) = (3, 4) gives the smallest possible values larger than that of the wildtype particle (Fig. 5C, dashed line). This angle is probably the most favored for the contact surface of cpAaLS(119) pentamers. Meanwhile, these mathematical analyses suggest the possibility of other pentagon-based tubular assemblies by modulating the contact surface or bending angles between capsomers.

The seemingly distinct structure of the cpAaLS twisted tubes is geometrically related to that of the corresponding straight tubes. This is illustrated by the interaction network analysis that can predict alternatives by rewiring the connections of a given structure (*46*). Applying this approach, the cpAaLS straight tube features squashed hexagons drawn by connecting the centers of interacting pentamers (Fig. 5D, black and blue solid lines). The only alternative to this can be obtained by deleting two existing contacts (blue solid lines) while generating two new ones (red dashed lines), yielding a triangle-based network that corresponds to the tiling blueprint of the twisted tubes (Fig. 5E). This geometrical similarity could explain why a subtle change in the amino acid sequence and assembly condition resulted in the transformation between straight and twisted tubes. Furthermore, the network rewiring analysis implies that no other tiling patterns exist with this type of pentamer.

## Conclusion

Successful control over the assemblies and their near-atomic resolution structures revealed the molecular origin of the dynamic, polymorphic nature of a circularly permuted cage-forming protein. A short peptide domain, which is untethered from the native position by topological rearrangement, inhibits the 3-fold symmetry interaction and holds the building blocks with a certain torsion angle, leading to the dramatic conversion of the wild-type dodecahedron to the previously unknown helical arrangements of pentameric subunits. Considering another cpAaLS variant, NC-4, adopts a quasi-equivalent assembly consisting of pentamers and hexamers (*43*), these results highlight the morphological plasticity of AaLS and circular permutation as a powerful approach for modulating or potentially customizing cage-like structures. The specific tiling patterns observed for the tubular structures are the most optimal blueprints for this particular pentamer. Meanwhile, geometrically feasible structures are more diverse, suggesting the potential existence of other types of assemblies formed by naturally occurring or engineered proteins. Notably, the mathematical approaches used in this study, AngelaR, the helix builder, the planar lattice model, and the interaction network analysis (*46*), are general and should be, therefore, useful for characterizing or predicting yet unknown polygon-based structures. While modular and readily modifiable cpAaLS(119) tubes are themselves an attractive platform for the prospective nanodevice development (*16*), our molecular and mathematical insights into the unique pentamer-based nanoarchitectures afford guidelines for the design of protein nanotubes with customized morphology and assembly dynamics.

## Acknowledgments

This publication was partially developed under the provision of the Polish Ministry and Higher Education project Support for research and development with the use of research infrastructure of the National Synchrotron Radiation Centre SOLARIS under contract nr 1/SOL/2021/2. We acknowledge Dr. Michał Rawski, Dr. Paulina Indyka, Dr. Marcin Jaciuk, Dr. Artur Biela, Grzegorz Ważny, as well as the Structural Biology Core Facility at the Malopolska Centre of Biotechnology of Jagiellonian University (MCB, JU) (supported by the TEAM TECH CORE FACILITY/2017-4/6 grant from the Foundation for Polish Science) for their support in cryo-EM experiments. We are also grateful for the Polish high-performance computing infrastructure PLGrid (HPC Centers: ACK Cyfronet AGH) for providing computer facilities and support within computational grant no. PLG/2020/014009. We thank Dr. Jakub Nowak (MCB, JU) and Dr. Neil A Ranson (University of Leeds) for their help in Nanotemper DSF/DLS experiments and cryo-EM helical reconstruction, respectively. The open-access publication of this article is funded by the Priority Research Area BioS under the program “Initiative of Excellence—Research University” at JU.

## Author contributions

Conceptualization: YA

Methodology: LK, FF, MB, RT, YA

Investigation: LK, FF, MA, JGH, RT

Visualization: LK, FF

Funding acquisition: RT, YA

Project administration: YA

Supervision: RT, YA

Writing – original draft: LK, FF, RT, YA

Writing – review & editing: LK, FF, JGH, RT, YA

## Data and materials availability

The principal data supporting the findings of this study are provided in the figures and supplementary materials. The code implementing the mathematical analysis is available from GitHub: https://github.com/MathematicalComputationalVirology/TubeModeler. Cryo-EM maps and atomic models have been deposited in the Electron Microscopy Data Bank (EMDB) and the Worldwide Protein Data Bank (wwPDB), respectively, with the following accession codes: EMDB-51006 and PDB 9G3P (12-pentamer cage), EMDB-51005 and PDB 9G3O (24-pentamer cage), EMDB-51004 and PDB 9G3N (36-pentamer cage), EMDB-51003 and PDB 9G3M (straight tube), EMDB-51001 and PDB 9G3J (28-Å gap twisted tube), EMDB-51000 and PDB 9G3I (18-Å gap twisted tube), and EMDB-50999 and PDB 9G3H (0-Å gap twisted tube).

## Supplementary Materials

Materials and Methods

Figs. S1 to S16

Tables S1 to S3

