## Supplementary Materials for "Dynamic assembly of pentamer-based protein nanotubes"

**The PDF file includes:**

Materials and Methods
Figs. S1 to S16
Tables S1 to S3

### Materials and Methods

Materials

All chemicals and biochemicals were purchased from Sigma-Aldrich (Burlington, MA, USA), New England BioLabs (Ipswich, MA, USA), or Thermo Fisher Scientific (Waltham, MA USA). Oligonucleotides were synthesized by Sigma-Aldrich. *E. coli* strains BL21-Gold(DE3) and DH5α competent cells were purchased from Agilent (Santa Clara, CA, USA) and Thermo Fisher Scientific, respectively. The plasmid pMG_cpAaLS_L8(119) (*39*) and pMG_cpAaLS_L8(84) (*41*) were kindly provided by Prof. Donald Hilvert (ETH Zurich, Switzerland).

Molecular cloning

The primers and plasmids used in this study are listed in Table S1 and S2, respectively. Plasmid pMG_cpAaLS_L8(119Δ120-131) was prepared from pMG_cpAaLS_L8(119) by cassette cloning via NdeI and XhoI sites. Plasmid pMG_cpAaLS_L8(119, C37S) was prepared from pMG_cpAaLS_L8(119) by site-directed mutagenesis. Plasmid pMG_cpAaLS_L8(119, C37S) was used to prepare pMG_cpAaLS_L8(119, C37S, A85C) by site-directed mutagenesis. *E. coli* strain DH5α was used as the host cells for every cloning step. Sequences of plasmids were confirmed by DNA Sanger sequencing performed by Eurofins Genomics Europe Sequencing GmbH (München, Germany).

Protein expression

All proteins were produced in *E. coli* strain BL21-Gold(DE3) transformed with pMG vectors. The cells were cultured at 37 °C and 220 rpm in 0.5 L of lysogeny broth, Miller formulation (LB) medium supplemented with 100 µg/ml ampicillin until the OD600 reached ~0.6, at which point protein production was induced by the addition of 0.5 mM isopropyl β-D-1-thiogalactopyranoside (IPTG). After culturing at 25 °C and 180 rpm for 20 hrs, cells were harvested by centrifugation at 3,000 ×g, 4 °C for 10 min and then stored at –20 °C until protein purification.

Protein purification

Cell pellets from 250-mL cultures were resuspended in 100 mL of lysis buffer (25 mM Tris-HCl buffer (pH 8.5) containing 200 mM NaCl, 1 mM EDTA, and 1.2 mM MgOAc) supplemented with lysozyme (0.5 mg/mL) and DNaseI (5 µg/mL). After cell lysis by sonication, the insoluble fraction was removed by centrifugation for 25 min at 9,500 ×g and 25 °C. The supernatant was heated at 70 °C for 1 hr while stirring, followed by removal of insoluble fractions by centrifugation for 25 min at 9,500 ×g and 25 °C. The supernatant was then diluted in a 1:1 volume ratio with 50 mM Tris-HCl buffer (pH 8.5) and loaded onto anion exchange HiTrap Q HP columns (4× 5 mL, Cytiva). After washing with 50 mM Tris-HCl buffer (pH 8.5) containing 100 mM NaCl, protein was eluted using a 0.1-1 M NaCl gradient. Fractions at approximately 400 mM NaCl were pooled and NaCl concentration was reduced to <2 mM with 50 mM Tris-HCl buffer (pH 8.5) using an ultrafiltration centrifugal filter unit (30 MWCO, Amicon Ultra-15, Merck Millipore). The protein solution was loaded onto an anion exchange Mono Q 5/50 GL column (Cytiva). After washing with the 50 mM Tris-HCl buffer (pH 8.5) containing 100 mM NaCl, the protein sample was eluted with a 0-1 M NaCl gradient. Fractions at approximately 400 mM NaCl were pooled and NaCl concentration was reduced to <2 mM with 5 mM Tris-HCl buffer (pH 8.5) using ultrafiltration. The protein sample was concentrated to approximately 300 µM and subjected to size-exclusion chromatography (SEC) using a Superdex 200 increase 10/300 column (Cytiva) running with 5 mM Tris-HCl buffer (pH 8.5) at room temperature (RT) and a flow rate of 0.75 mL/min. Peak fractions were pooled and protein was concentrated to approximately 345 µM (with respect to monomer concentration unless specified herein after) and kept at RT until further experiments. Protein concentration was routinely determined by absorbance at 280 nm ($\varepsilon_{280}$ = 13,980 $M^{-1}$ $cm^{-1}$). Protein purity at each purification step was confirmed by SDS-PAGE with Coomassie R350 staining (fig. S16).

<u>Molecular mass analysis of cpAaLS(119) capsomers</u>

Apparent molecular mass of cpAaLS(119) in 50 mM Tris-HCl buffer (pH 8.5) was estimated by SEC coupled with right- and low-angle light scattering (RALS/LALS) detections (OMNISEC REVEAL, Malvern Panalytical Ltd, Malvern, UK) as described previously (*11*). Protein sample was buffer exchanged with 50 mM Tris-HCl buffer (pH 8.5) to 345 µM using ultrafiltration and analyzed by SEC-RALS/LALS using a Superdex 200 increase 10/300 column running at RT and 0.5 mL/min. Data was analyzed using Omnisec software (v11.10.7248.3) and parameters as follows: dn/dc = 0.185; dA/dC = 0; second virial coefficient (A2) = 0; RI = 1.333, viscosity = 0.7148 mPaS; detector oven temperature = 25°C. The system was calibrated with conalbumin (75 kDa, 345 µM, 500 µL) (Gel Filtration High Molecular Weight Calibration Kit, GE Healthcare) in 50 mM Tris-HCl buffer (pH 8.5). Parameters used for the standard were as follows: intrinsic viscosity (IV, dL/g) = 0; the ratio between weight- and number-averaged molecular weight (MW/MN) = 1; a = 0.7 (Mark-Houwink parameter a). Settings for calculation method used for assessing molecular weight included: calibration type – triple detection, analysis type – calculate sample concentration from dn/dc.

<u>Assembly of cpAaLS(119) protein cages</u>

Protein sample in 5 mM Tris-HCl buffer (pH 8.5) was mixed in 1:1 volume ratio with 2× buffer, followed by ultrafiltration in a corresponding 1× buffer [50 mM Tris-HCl buffer (pH 7.5, 8.0, or 8.5) containing 0 mM, 150 mM, 350 mM, or 1 M NaCl]. Protein was concentrated to approximately 172 µM and kept at RT for approximately 16 hrs, and then subjected to SEC using a Superose 6 increase column (Cytiva) at RT and a flow rate of 1 mL/min. The chromatograms were analyzed with ChromLab software (Bio-Rad) (fig. S2B-E, left). Prior to SEC, each sample was analyzed by cryo-EM using a Glacios microscope (Thermo Fisher Scientific).

<u>Cryo-EM grid preparation</u>

Approximately 3.5 µL of the sample was applied onto a freshly glow discharged TEM grid (Quantifoil R2/2, Cu 200 mesh) and plunge vitrified into liquid ethane by a Vitrobot Mark IV (Thermo Fisher Scientific). For cryo-EM with a Glacios microscope, the following vitrification parameters were used: humidity 95%, temperature 10 °C. blot total 1, wait time 30 sec, and drain time 0 sec. Blot force and blot time were adjusted depending on the ionic strength of the buffer; 0 mM NaCl: 7 and 7 sec, 150-350 mM NaCl: 6 and 6 sec, 1 M NaCl: 5 and 5 sec, respectively. For cryo-EM measurement on a Titan Krios G3i microscope (Thermo Fisher Scientific), the following vitrification parameters were used: humidity 95%, temperature 10 °C. blot total 2, wait time 30 sec, blot force 3, blot time 3 sec, and drain time 0 sec.

Cryo-EM with Glacios microscope

All the cryo-EM micrographs were collected at the Cryo-EM Centre of the National Synchrotron Radiation Centre SOLARIS (Krakow, Poland). The micrographs, typically 20 for each variant and condition, were acquired on a Glacios microscope (Thermo Fisher Scientific) fitted with a Falcon 4 detector operated at 200 kV accelerating voltage, magnification of ×150k, and corresponding pixel size of 0.96 Å/px. The collected micrographs were analyzed using Fiji (*50*) and cryoSPARC v4.2.1 (*51*).

Disassembly of cpAaLS(119) protein cages

Protein solution in 5 mM Tris-HCl buffer (pH 8.5) was mixed in 1:1 volume ratio with 2× buffer, followed by ultrafiltration in a corresponding 1× buffer [50 mM Tris-HCl buffer (pH 8.0) containing 0.15 or 1 M NaCl] (fig. S3A). Protein was concentrated to approximately 172 µM and kept at RT for approximately 16 hrs. The protein assemblies were analyzed and isolated by SEC using a Superose 6 increase 10/300 column in corresponding 1× buffer at RT and a flow rate of 1 mL/min. Peak fractions were pooled, and protein was concentrated to approximately 90 µM in 50 mM Tris-HCl buffer (pH 8.5). After approximately 1 hr, the sample was reanalyzed by SEC using the same column and conditions, except for the eluent of 50 mM Tris-HCl buffer (pH 8.5).

Thermal stability of cpAaLS assemblies

Protein sample in 5 mM Tris-HCl buffer (pH 8.5) was mixed in 1:1 volume ratio with 2× buffer, followed by ultrafiltration in a corresponding 1× buffer [50 mM Tris-HCl buffer (pH 7.5) containing 0.15 or 1 M NaCl; or 50 mM Tris-HCl buffer (pH 8.5) containing 0 or 1 M NaCl] (fig. S4A). Protein was concentrated to approximately 172 µM and kept at RT for approximately 16 hrs. The protein solution was diluted to 57 µM with the corresponding buffers and transferred to a Prometheus high sensitivity glass capillary sealed with a dedicated sealing paste (NanoTemper Technologies). Thermal shift assay with differential scanning fluorimetry (DSF) and dynamic light scattering (DLS) detections were performed using a Prometheus PANTA instrument (NanoTemper Technologies) over a 25-110 ºC temperature range with a 1 ºC/min ramp. The measurements were triplicate. Datasets were analyzed and merged using the PR.PantaAnalysis software.

Cryo-EM single particle reconstruction of cpAaLS(119) capsomers

Cryo-EM data were acquired on a Titan Krios G3i microscope operated at 300 kV accelerating voltage, magnification of ×105k, and pixel size of 0.86 Å/px. A K3 direct electron detector used for data collection was fitted with BioQuantum Imaging Filter (Gatan) using a 20 eV slit and operated in counting mode. Imaged areas were exposed to 40 $e^-/Å^2$ total dose (corresponding to ~16 $e^-$/px/s dose rate measured in vacuum). Forty-frame movie stacks were obtained using under-focus optical conditions with a defocus range of −2.1 to −0.9 µm and 0.3 µm steps. The collected datasets were analyzed using cryoSPARC v4.2.1. First, "Patch Motion Correction" and "Patch CTF Estimation" steps were performed. Next, a "Blob Picking" step resulted in 224767 particles, and subsequent 2D classification produced classes corresponding to flat pentamers and pentamer pairs. Due to the preferential orientation of particles in the grid holes, 3D reconstruction was unsuccessful.

Cryo-EM single particle reconstruction of cpAaLS spherical cages

The cpAaLS(84) 16-nm cages, cpAaLS(119) 24-nm or 36-nm spherical assemblies were isolated by SEC using a Superose 6 increase 10/300 column in 50 mM Tris-HCl buffer (pH 8.0) containing 150 mM, 500 mM or 350 mM NaCl. The peak fractions were concentrated by ultrafiltration to approximately 28 µM (or 56 µM for cpAaLS(119) 24-nm cages) in the corresponding buffer and used for vitrification. To reduce salt concentration, the sample with cpAaLS(119) 24-pentamer cage was subjected to a quick 2-fold dilution with 50 mM Tris-HCl buffer (pH 8.0) (final 250 mM NaCl) prior to vitrification. Cryo-EM data was collected on a Titan Krios microscope, as described above with minor modifications, and analyzed using RELION v3.1 (*52*) using parameters shown in fig. S5. Briefly, after motion correction and CTF estimation, approximately 500 particles were picked manually, and used for 2D classification as well as the generation of preliminary classes for template picking. After Ab initio reconstruction in C1 using picked particles, 3D classification and 3D refinements were performed using icosahedral (I) or tetrahedral (T) symmetries (fig. S5). The particle stacks were subjected to iterative per-particle defocus and global CTF refinements, followed by Bayesian polishing. Gold-standard Fourier shell correlation and local map resolutions were calculated with 0.143 FSC cut-off. Prior to model fitting, the combined half-maps were sharpened with DeepEMhancer.

Cryo-EM helical reconstruction of cpAaLS(119) straight tube

As for spherical cages, the cpAaLS(119) straight tube fraction was isolated by SEC using a Superose 6 increase 10/300 column in 50 mM Tris-HCl buffer (pH 8.0) containing 150 mM NaCl and concentrated by ultrafiltration to approximately 28 µM. Cryo-EM data were collected on a Titan Krios microscope, as described above, using a magnification of ×81k and a pixel size of 1.1 Å/px. Forty-frame movie stacks were obtained using super-resolution mode (0.55 Å/px) and under-focus optical conditions with a defocus range of −3.0 to −0.9 µm and 0.3 µm steps. The collected dataset was analyzed using helical reconstruction in RELION v3.1 (*53*) using parameters shown in fig. S6. Briefly, after motion correction (2× bin) and CTF estimation, straight helical segments were manually picked by selecting start-end coordinates and subjected to 2D classification. A cylinder with a 210-Å outer diameter was generated using relion_helix_toolbox and used as an initial model for 3D classification. The final 168323 particle stacks were used for 3D helical reconstruction, followed by iterative per-particle defocus, global CTF refinements, and Bayesian polishing. Gold-standard Fourier shell correlation and local map resolutions were calculated with 0.143 FSC cut-off. Prior to model fitting, the combined half-maps were sharpened with DeepEMhancer (*54*).

Assembly of cpAaLS(119, C37S, A85C) twisted tube

The cpAaLS(119, C37S, A85C) variant in 5 mM Tris-HCl buffer (pH 8.5) was mixed with 50 mM Tris-HCl buffer (pH 7.0) followed by buffer exchange using ultrafiltration with the same buffer (fig. S7). Protein was concentrated to approximately 172 µM and kept at RT for approximately 16 hrs. The resulting solution was then mixed in 1:1 volume ratio with 50 mM Tris-HCl buffer (pH 8.5) containing 0.3 M NaCl, followed by ultrafiltration in 50 mM Tris-HCl buffer (pH 8.5) containing 0.15 M NaCl and 1 mM tris(2-carboxyethyl)phosphine (TCEP). Protein was concentrated to approximately 172 µM and kept at RT for approximately 16 hrs. The protein assembly was then analyzed and isolated by SEC using a Superose 6 increase 10/300 column in 50 mM Tris-HCl buffer (pH 8.5) containing 0.15 M NaCl and 1 mM TCEP at RT and a flow rate of 1 mL/min. Peak fractions corresponding to nanotubes were pooled and protein was concentrated to approximately 57 µM for cryo-EM analysis. The analog experiment was performed with the parent cpAaLS(119) variant for comparison.

Cryo-EM helical reconstruction of cpAaLS(119, C37S, A85C) twisted tube

Cryo-EM data was collected on a Titan Krios G3i, as described above, using a magnification of 105k and a corresponding pixel size of 0.846 Å/px. Movie stacks (40 frames) were obtained using under-focus optical conditions with a defocus range of -1.5 to -0.9 µm and 0.3 µm steps. The collected dataset was analyzed using 'Helical Reconstruction' in cryoSPARC v4.4.1 (*51*). First, "Patch Motion Correction" and "Patch CTF Estimation" steps were performed. Next, approximately 500 particles were picked manually. The acquired particles were subjected to 2D classification and used in the generation of preliminary classes for the subsequent template picking using the filament tracing tool (fig. S8). Particles were extracted using 2× binning. Following 2D classification, a cylinder with a 200/130 Å outer/inner diameter was generated and used as an initial model for initial 3D "Helical Refinement". Following "Heterogeneous Refinements", the particles sets were split based on uniformity into three independent structures with 1) ~24.5-, 2) ~22-, or 3) ~20-Å helical rise. The resulting structures correspond to "twisted tube" with 1) ~28-Å, 2) 18-Å, or 3) no/0-Å gap between dual helical threads. Prior to final 3D helical refinements, the particles were unbinned and subjected to additional 2D classification resulting in corresponding 192310, 807640, or 208379 stacks. Furthermore, the particles were subjected to "Reference Based Motion Correction". During final 3D "Helical Refinements", the particles and micrographs were subjected to per-particle defocus, global CTF refinements, and Ewald Sphere correction to generate high-resolution maps. Gold-standard Fourier shell correlation and local map resolutions were calculated with 0.143 FSC cut-off. Prior to model fitting, the combined half-maps were sharpened with DeepEMhancer (*54*).

Molecular modeling

The initial atomic model was sourced from a X-ray crystal structure (1.6-Å resolution) of wild-type AaLS (PDB: 1HQK) (*35*). Following rigid body fitting using ChimeraX v1.7 (*55*), and manual modification in Coot (*56*), coordinates were flexibly fit with Isolde . The models were real-space refined in Phenix v1.20.1-4487. The final coordinates were validated using MolProbity (*57*) and the model statistics are presented in Table S3. The cryo-EM maps and atomic models were displayed using ChimeraX.

Estimation of the bending and torsion angles

The normal vectors to the planes defined by two adjacent pentamers were computed. If these vectors are in a plane with a line connecting the centers of the pentamers, the torsion angle is zero and the bending angle is 180 minus the angle between two normal vectors, which corresponds to the bending angle at the interface of the pentamers. If the normal vectors do not form a plane with the line connecting the centers of the pentamers, the torsion angle corresponds to the angle between the normal vector before and after it has been rotated into that plane. The code implementing this procedure is available from GitHub:
(https://github.com/MathematicalComputationalVirology/TubeModeler "Script_measuring_angles.py").

Simulation of the single-stranded helix model for twisted tubes

The helix model was constructed by applying bending and torsion angles iteratively starting with an initial pentamer. The incoming pentamer is located at a position on the interface specified by a free parameter in the code. At that point, the construction is fully determined. The code implementing this procedure is available from GitHub:

(https://github.com/MathematicalComputationalVirology/TubeModeler "Twisted_Tube_3D.m").

Mathematical characterization of the straight tubes

The mathematically viable straight tubes with helicity number $n_h$ and periodicity number $n_p$ are characterized based on the construction shown in SI Fig. S2. Taking the center of the green pentagon as the origin ($O$), its vertices $P_i$ correspond to the 5th roots of unity:

$$P_i = \left(\cos\left(\frac{2(i-1)\pi}{5} + \frac{\pi}{10}\right), \sin\left(\frac{2(i-1)\pi}{5} + \frac{\pi}{10}\right)\right), i = 1, 2, ..., 5.$$

Then the vector $\vec{H} = (a, b)$ is the translation vector between two strips. The vertices of the translated pentagon are $Q_i = P_i + \vec{H}$ for i = 1, 2, ..., 5. Denoting by $R_1$ the intersection point of the lines $\overline{P_1P_2}$ and $\overline{Q_1Q_5}$ and using coordinates values, one obtains:

$$R_1 = \left(-\frac{b-ac}{\sqrt{10+2\sqrt{5}}} + \frac{\sqrt{10+2\sqrt{5}}}{4}, \frac{\sqrt{10-2\sqrt{5}}(b-ac)}{(\sqrt{5}+1)\sqrt{10+2\sqrt{5}}} + \frac{\sqrt{5}-1}{4}\right),$$

where

$$c = \frac{\sqrt{10+2\sqrt{5}}}{\sqrt{5}-1}.$$

We choose the point $R_1'$ on $P_3P_4$ such that the length of $P_4R_1'$ equals that of $R_1P_2$. As the contact pattern is the same along a strip, the length of $P_4R_1'$ moreover equals that of $TR_3$. Thus, $\vec{P} = \overrightarrow{R_1'R_3} = R_3 - R_1'$ and $R_3 = R_1 + (P_1 - P_3)$. Since $P_4R_1' = R_1P_2$, $R_1'$ is an anticlockwise rotation by 144° of $R_1$ around the origin, i.e.,

$$R_1' = \begin{bmatrix} \cos(144) & -\sin(144) \\ \sin(144) & \cos(144) \end{bmatrix} R_1.$$

To obtain the helicity number $n_h$ and periodicity number $n_p$, the following equality should be satisfied for the second components of the vectors $\vec{H}$ and $\vec{P}$:

$$n_h \vec{H}(2) = n_p \vec{P}(2).$$

By substitution, the above equality then leads to the following relation:

$$a = \frac{n_p - 2n_h}{cn_p} b.$$

This equation demonstrates that the parameters $n_h$ and $n_p$ are geometrically linked.

Procedure for mathematical model building of straight tubes

The values of $b$, $n_p$ and $n_h$ define $a$, which gives vectors $\vec{H}$ and $\vec{P}$. Starting with a pentagon, and using $b = 2.18$ to match the biological structure, the coordinates of the pentagon are translated as shown in Fig. S11. The contact area corresponds to the proportion of a pentagonal edge in contact with the edge of a neighboring pentamer, and it is given as a percentage, i.e., $100 \times \frac{P_1R_1}{P_1P_2}$. The code implementing this procedure is available from GitHub:

(https://github.com/MathematicalComputationalVirology/TubeModeler "Straight_Tube_2D.m" and "Straight_Tube_3D.m").

Rewiring of the capsomer network (*46*)

The alternative architectures were obtained from a given structure with reference to its interaction network. In the case of the straight tubes, the interaction network is given by squashed hexagons, which are composed of two triangles and one square. Associating weights (including zero) to these edges, respecting (helical) symmetry-equivalent positions and connectivity, results in different structures. The only other option here is the twisted tube architecture.

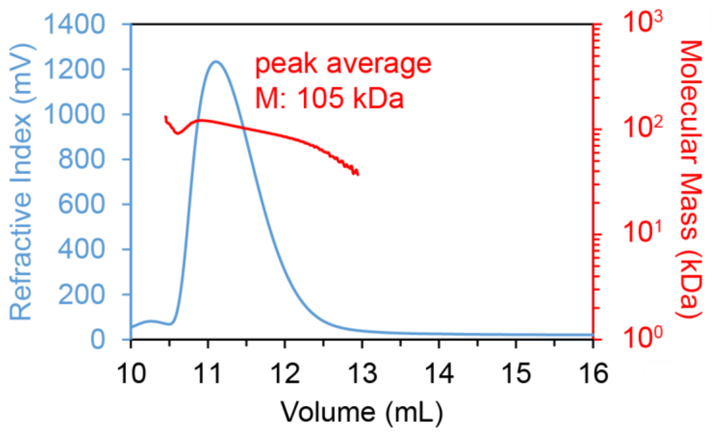


**Fig. S1. Disassembly of cpAaLS(119) cages in a low-ionic-strength buffer at alkaline pH.** The cpAaLS(119) protein was analyzed by size-exclusion chromatography coupled with right/low-angle light scattering detectors (SEC-RALS/LALS) in 50 mM Tris-HCl buffer at pH 8.5, estimating the average molecular mass (M) of 105.2 kDa. This likely corresponds to a mixture of cpAaLS pentamer (86.5 kDa) and its dimer (174.3 kDa), which was further confirmed by cryo-EM (Fig. 2A).

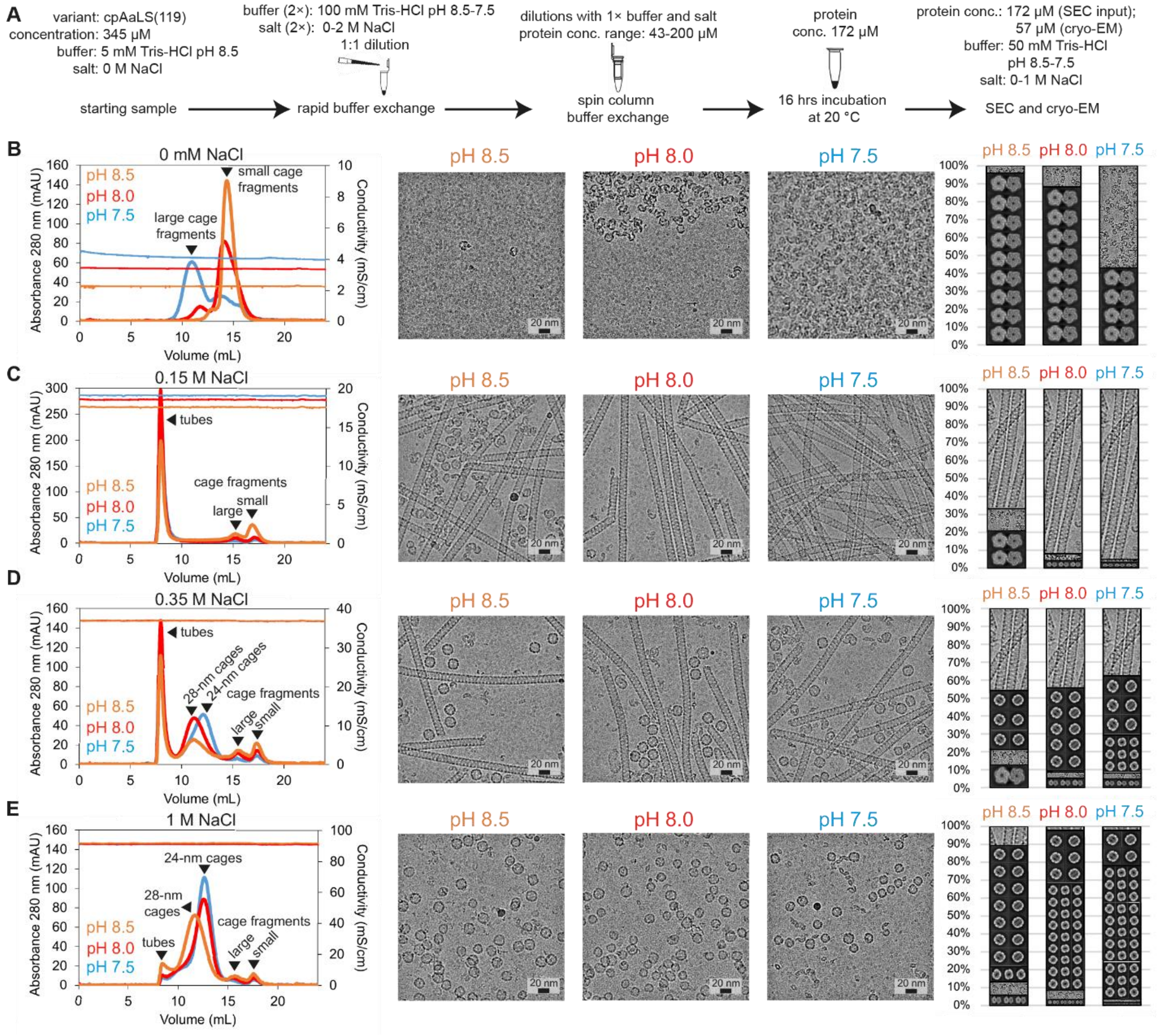


**Fig. S2. Ionic strength and pH-dependent assembly of cpAaLS(119).**

(A) Experimental scheme to assemble cpAaLS(119) proteins under different buffer conditions. (B-E) Size-exclusion chromatography (SEC) profiles (left), representative cryo-EM micrographs (middle), and relative distribution of cpAaLS(119) assembly states (right) in a buffer containing 0 (B), 0.15 (C), 0.35 (D), or 1 M (E) NaCl at pH 8.5 (orange), 8.0 (red), or 7.5 (blue). The relative distribution of the assembly states was quantified by the integrating peak areas of the SEC traces. The 24-nm and 28-nm cages were discriminated by manual inspection and classification of spherical particles from ~20 micrographs. Small and large cage fragments in SEC profiles likely represent different oligomerization statuses of pentamers. In the SEC profiles, conductivities are also shown as a reference for the buffer ionic strength which is slightly dependent on pH due to the different ionization levels of Tris base.

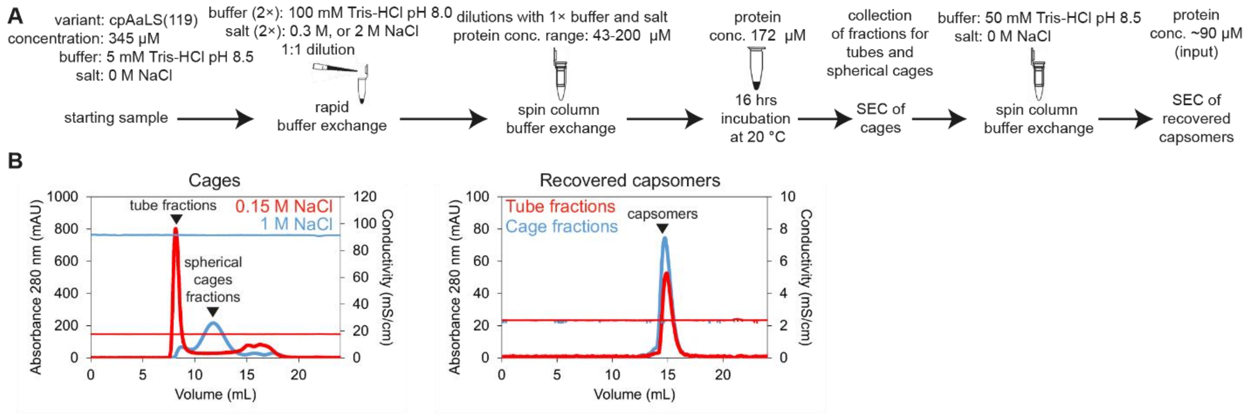


**Fig. S3. Salt and pH-dependent disassembly of cpAaLS(119) protein cages into capsomers.** (A) Experimental scheme describing stepwise cage assembly-disassembly by buffer exchange. (B) 50 mM Tris-HCl buffers (pH 8.0) containing 0.15 M (red) or 1 M (blue) NaCl were used for tubular and spherical cage formation, respectively, and the major assemblies were isolated by SEC (bottom left). Following buffer exchange to 50 mM Tris-HCl buffer (pH 8.5) resulted in nearly 100% cage disassembly into the fragments (capsomers, bottom right). Conductivities are shown as a reference for the buffer ionic strength.

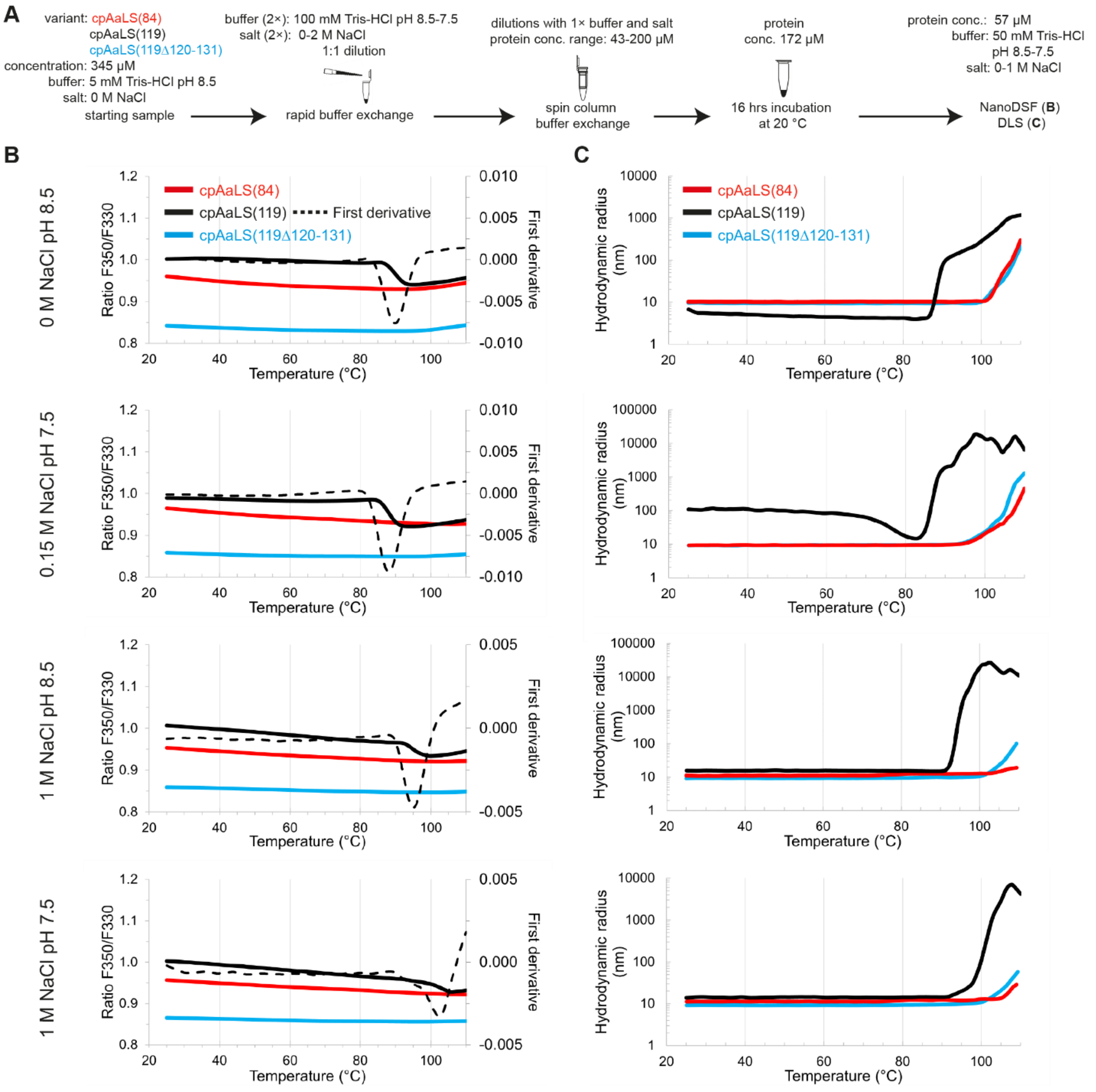


**Fig. S4. Thermal stability of cpAaLS assemblies.**

(A) Experimental scheme for preparation of cpAaLS(84) (red), cpAaLS(119) (black), and cpAaLS(119Δ120-131) (blue) samples in a varied NaCl concentration (0, 0.15, or 1 M) and pH (8.5, or 7.5). (B) Nano-differential scanning fluorimetry (NanoDSF) thermographs of these samples, in the temperature ramp 25–110 ºC. The melting temperature ($T_m$) of cpAaLS(119) was estimated by the first derivative (dashed lines) minima of the observed fluorescent ratio at F350/F330 to be 90, 88, 96, and 103 °C in the buffers where the major assembly status of the protein should be capsomers, tubular, 24-nm, and 28-nm cages, respectively. (C) Dynamic light scattering of the corresponding samples. Increasing hydrodynamic radii in the temperature ramp indicate aggregation of cpAaLS(119), ahead of cpAaLS(84) or cpAaLS(119Δ120-131).

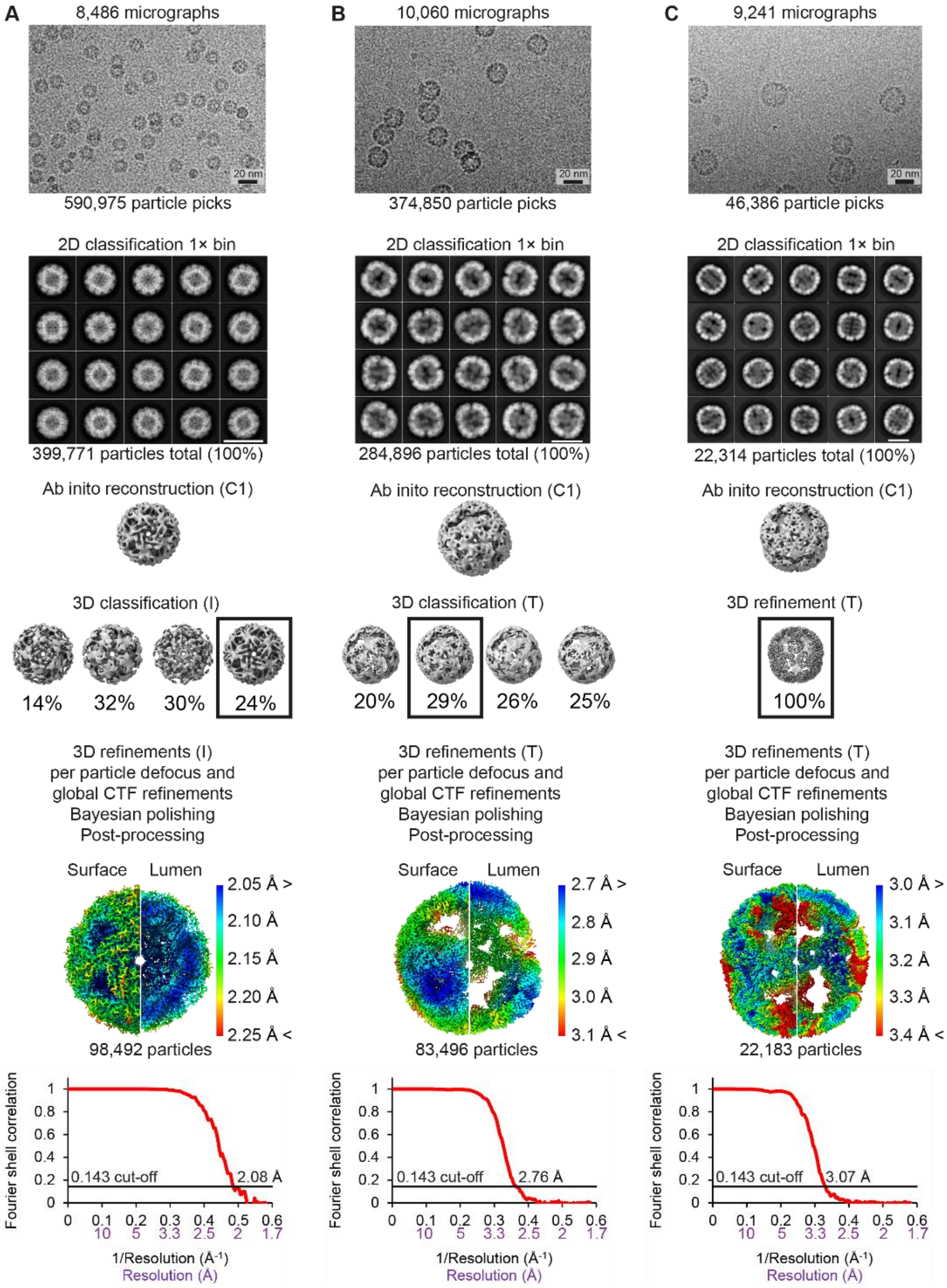


**Fig. S5. Single particle reconstruction of cpAaLS(84) and cpAaLS(119) spherical cages.** (A-C) The scheme represents the processing of cryo-EM datasets for the icosahedral (I) cpAaLS(84) 12-pentamer cage (A), tetrahedral (T) cpAaLS(119) 24- (B), or 36-pentamer (C) cages. From top to bottom, the scheme shows the representative micrographs (20 nm black scale bar), selected 2D classes (white scale bar), ab initio reconstruction, 3D classification with selected classes (boxed with black line), and lists per particle/micrograph/movie corrections. Following final 3D cryo-EM refinements, the maps were filtered and colored by local resolution (0.143 FSC cut-off). Gold-standard Fourier shell correlation curves are shown (bottom). The 3D maps are not to scale. The absolute and/or relative (% of total) number of particles is indicated after each selection step.

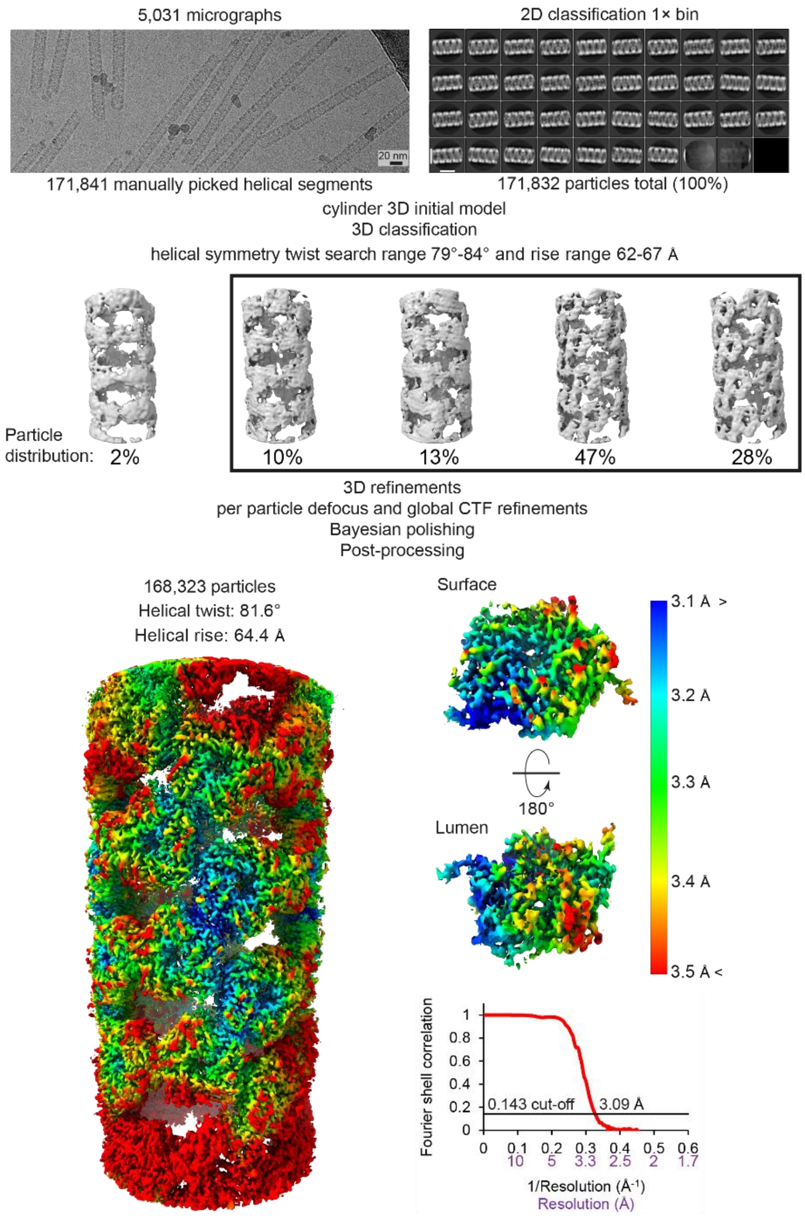


**Fig. S6. Helical reconstruction of cpAaLS(119) straight tube.**

The processing scheme of the cryo-EM dataset shows (left to right, top to bottom) a representative micrograph, selected 2D classes (white scale bar = 20 nm), 3D classification with selected classes (black box), and per particle/micrograph/movie corrections. Following final 3D cryo-EM refinements the map was filtered and colored by local resolution (0.143 FSC cut-off). The cryo-EM density corresponding to the asymmetric pentamer in the middle of the tubular segment was extracted to display the best local resolution on the surface and in the lumen of the cage. Gold-standard Fourier shell correlation curve was calculated with a 30% helical axis mask (right bottom). The 3D maps are not to scale. The absolute and/or relative (% of total) number of particles is indicated after each selection step.

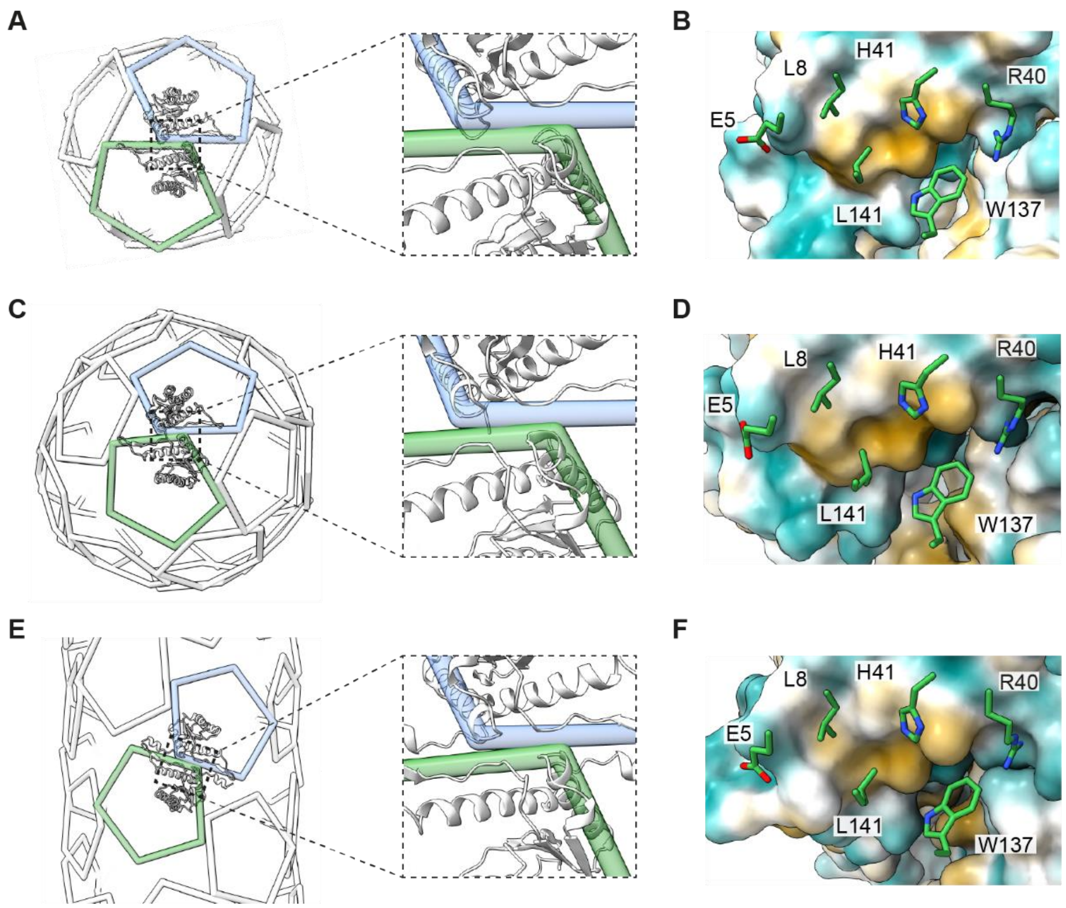


**Fig. S7. Bonding network between pentamers at the (pseudo) 2-fold symmetry in cpAaLS(119) cages.**

(A, C, E) Wire diagram of the 12-pentamer cpAaLS (84) (A), 24-pentamer cpAaLS(119) (C) spherical cages, and cpAaLS(119) straight tube assembly (E) with an enlarged view of a representative pentamer pair (green and blue), and cartoon representation of interfacing monomers. (B, D, F) The corresponding surfaces presenting hydrophobic (brown) and hydrophilic (blue) properties. The contacting residues from the opposite monomer are displayed as green sticks. Aside from the varied orientation of these side chains, all the assemblies preserve essentially the same amino acid interaction patterns at the (pseudo) 2-fold symmetrical interface.

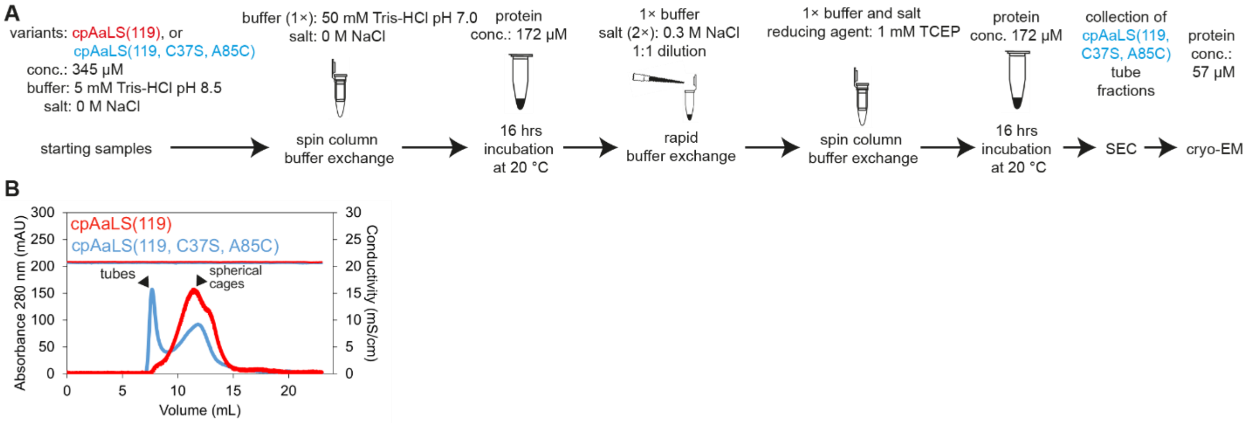


**Fig. S8. Assembly of cpAaLS(119, C37S, A85C) into twisted tubes.**

(A) Experimental scheme for preparation of the twisted tubes. (B) Size-exclusion chromatogram of the cpAaLS(119, C37S, A85C) sample (blue), showing a mixture of twisted tubes and spherical cages. The parent cpAaLS(119) variant forms spherical cages (red) almost exclusively under the same condition. Difference in the assembly states of cpAaLS(119) with 0.15 M NaCl from the results shown in fig S2C might be attributed to the incubation of the protein at pH 7.0, prior to addition of salt.

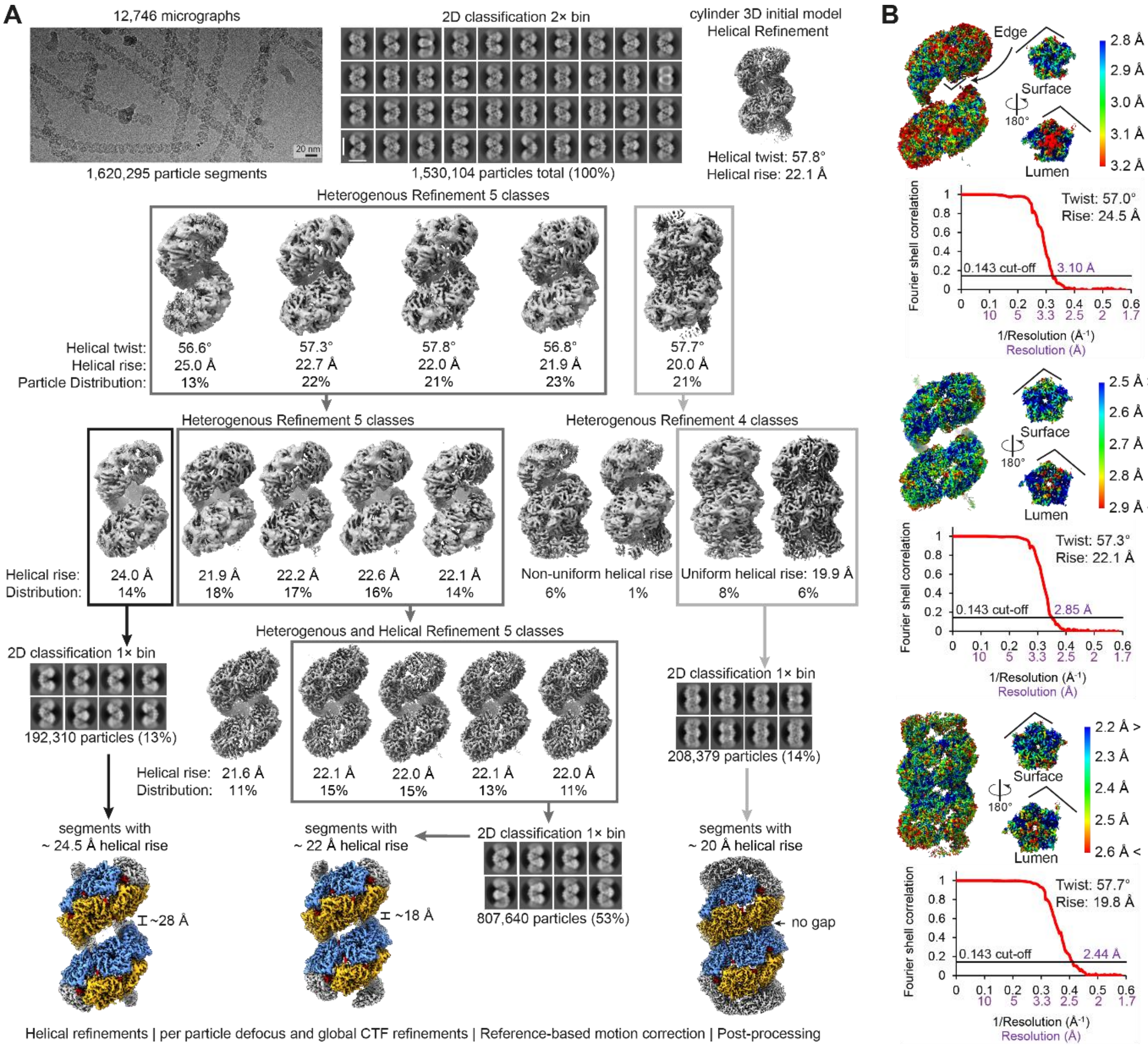


**Fig. S9. Helical reconstruction of cpAaLS(119, C37S, A85C) twisted tube.**

(A) Processing of the cryo-EM dataset showing the representative micrograph (20 nm black scale bar), selected 2D classes (20 nm white scale bar), and initial helical refinement. The non-uniform helical segments (particles) were separated based on the helical rise parameter. The segments with ~24.5-, ~22-, or ~20-Å helical rise were obtained by iterative 3D heterogeneous refinements (selected 3D classes are boxed with black, dark grey, and light grey lines, respectively), and confirmed by additional 2D classification. The final particle stacks were used to obtain uniform structures with ~28-Å, ~18-Å, or no gap (0-Å) between dual helical strips, respectively (orange/blue strips, bottom left, middle and right). The absolute and/or relative (% of total) number of particles is indicated after each selection step. (B) Following per particle/micrograph/movie corrections and final 3D cryo-EM refinements, the corresponding maps were filtered and colored by local resolution (0.143 FSC cut-off). The cryo-EM densities corresponding to representative asymmetric pentamers were extracted to display the local resolution on the surface and in the lumen of the cages. The edge of the pentamers exposed to the gap between helical strips is indicated. Gold-standard Fourier shell correlation curve is shown. The 3D maps are not to scale.

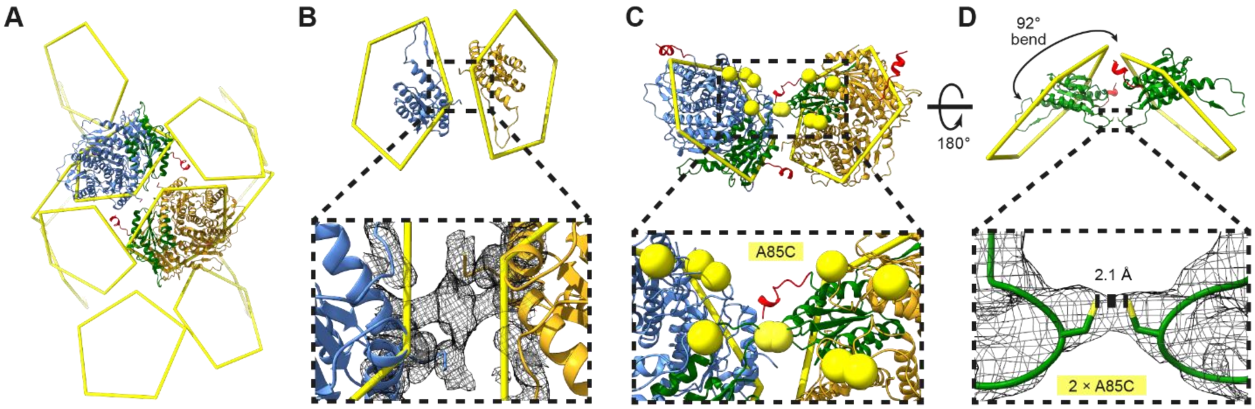


**Fig. S10. Potential disulfide bridge facilitating the acute bending angle in the pentamer-pentamer interaction.**

Potential disulfide bridge facilitating the acute bending angle in the pentamer-pentamer interaction. (A) The wire (yellow) representation of cpAaLS(119, C37S, A85C) twisted tube with no gap (0-Å gap) between dual strips. A pentamer pair interacting via acute bending angle is shown with ribbon representation. (B) Amplified view of the acute interaction between the adjacent protomers (blue and golden ribbons), indicating that this is not held by the hydrophobic (L8, L141, W137) or hydrophilic (E5, H41, R40) amino acids, unlike other pentamer-pentamer contacts exemplified in Fig. 2B, D, and F. The unfitted cryo-EM density (expanded region, black mesh) may correspond to the flexible 'untethered' α-helix(120-131). (C) Extracted pentamer pair shown from the lumen with A85C cysteines highlighted as yellow spheres. Two of the cysteines are in proximity (expanded region, green protomers). (D) The pentamer pair with isolated protomers (green ribbon) shown from the side. The continuous cryo-EM density (expanded region, black mesh) and optimal distance (2.1 Å) between the proximal A85C residues are suggestive of disulfide bond formation despite the presence of 1 mM TCEP. Similarly, close proximity of the A85C residues was also observed in 18- and 28-Å gap counterparts (not shown). Models and maps are not to scale.

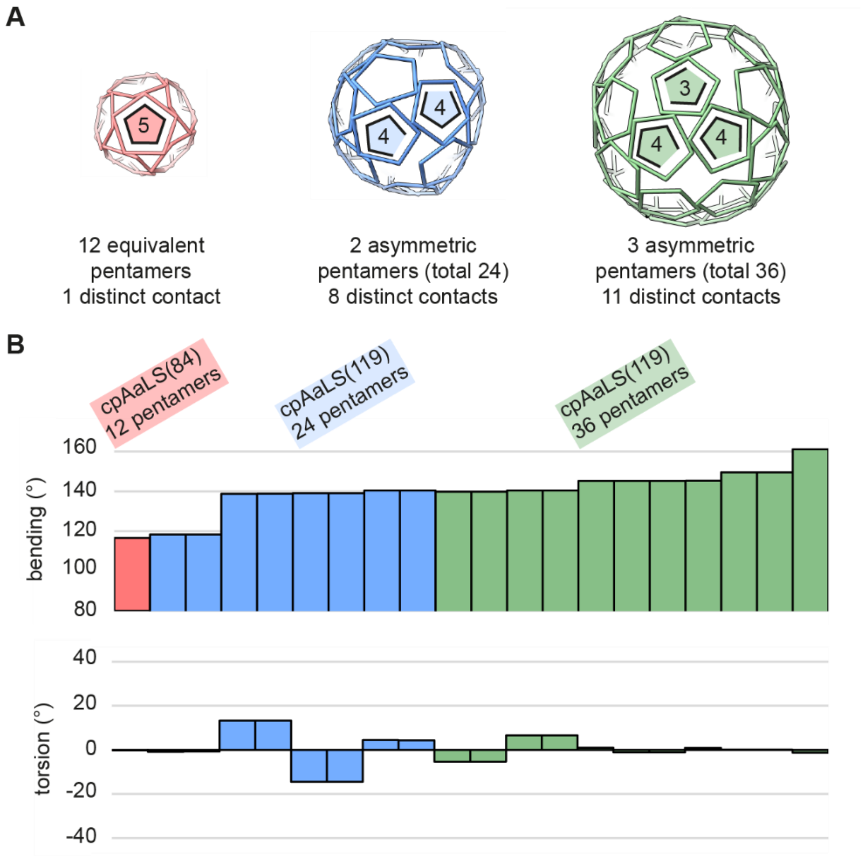


**Fig. S11. Capsomer interaction angles in spherical cages.**

(A) The wire representation of 12-pentamer cpAaLS(84) (red), 24-pentamer cpAaLS(119) (blue), and 36-pentamer cpAaLS(119) (green) assemblies. The numbers of contacts per each asymmetric pentamer are indicated. (B) Measured values of the bending (top) and the torsion angles (bottom) between two pentamers.

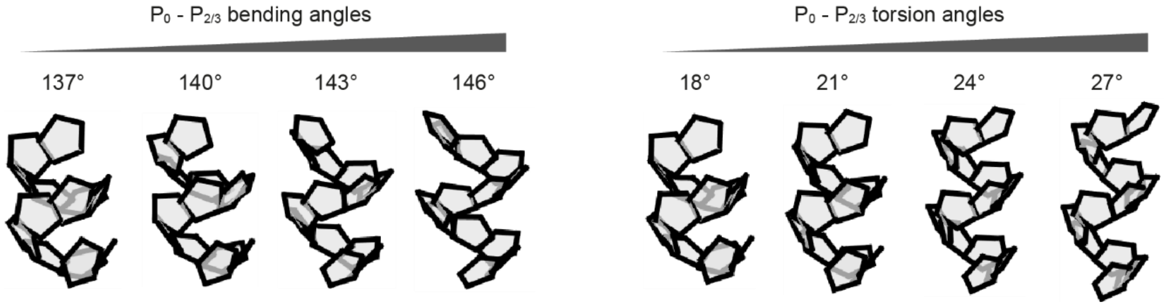


**Fig. S12. Correlation between pentamer interaction angles and helical rise.**

Computational simulation using a pentagon-based helical model showing that increasing bending (left) or torsion angle (right) for the $P_0$-$P_{2/3}$ pentamer interface (Fig. 4D) in twisted tubes results in the widening gap between helical strips. The simulation results agree with the experimental observations shown in Fig. 4, B to F.

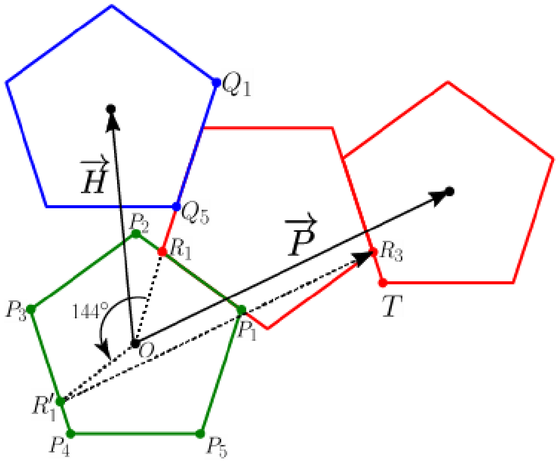


**Fig. S13. Mathematical characterization of the straight tubes.**

Taking the center of the green pentagon as the origin ($O$), vectors $\vec{H}$ and $\vec{P}$ specify the helicity and periodicity directions, respectively. Since the two adjacent green and red pentagons are translated as a unit, the origin is mapped to the center of the second pentagon (black dots). Labels refer to the variables used in the proof (Materials and Methods, Mathematical characterization of the straight tubes).

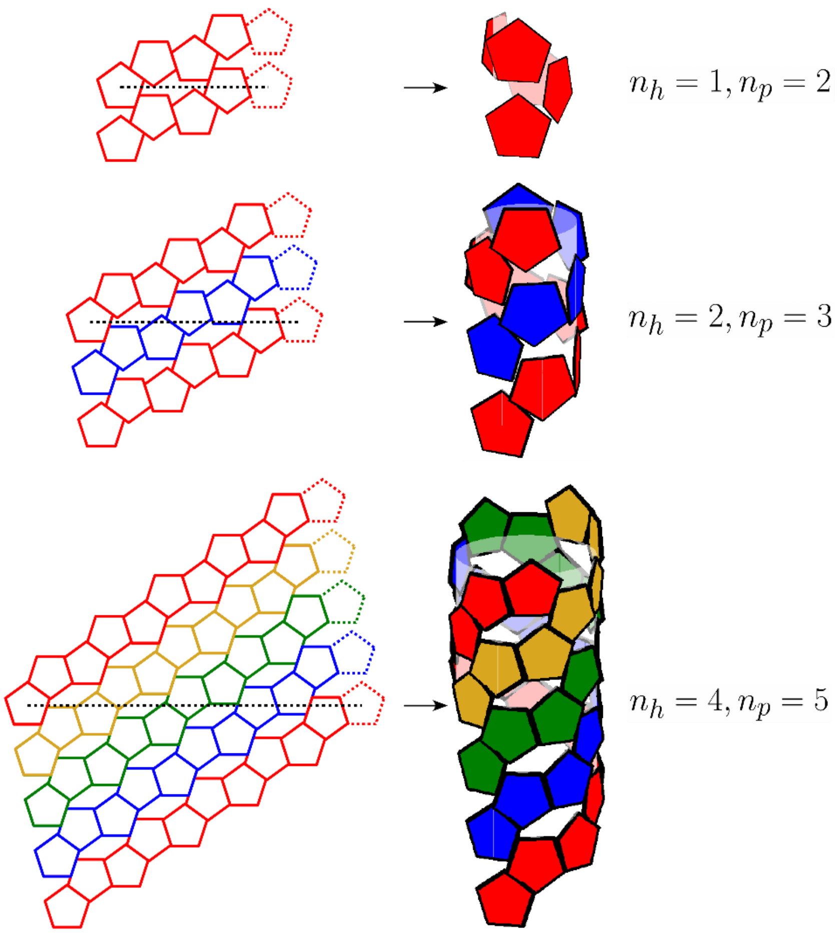


**Fig. S14. Tiling pattern and resulting tubular structures.**

Three geometrically possible tilling patterns with various $n_h$ and $n_p$ numbers and the corresponding 3D tube models are shown as examples. Individual pentamer threads are shown in different colors. The pentamers connected with the black dashed lines are identical in the 3D tubular structure.

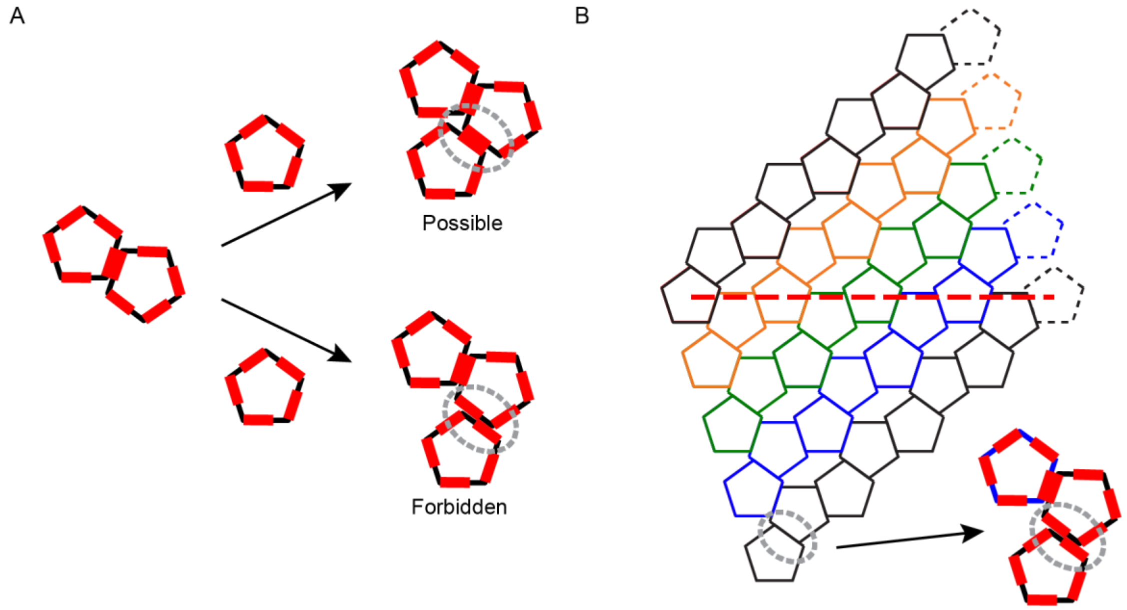


**Fig. S15. Tiling pattern violating the biologically occurring interactions.**

(A) Schematic representation of possible and forbidden interaction patterns for three pentamers having an asymmetric interaction surface (presented as red patches). (B) Example of a geometrically possible but biologically forbidden tilling pattern. This tilling pattern, defined by $(n_h, n_p)$ = (4, 4), requires flipped pentagons to retain the same type of interfaces, which is unlikely to occur with protein building blocks. The pentamers connected with the red dashed line are identical in the 3D tubular structure.

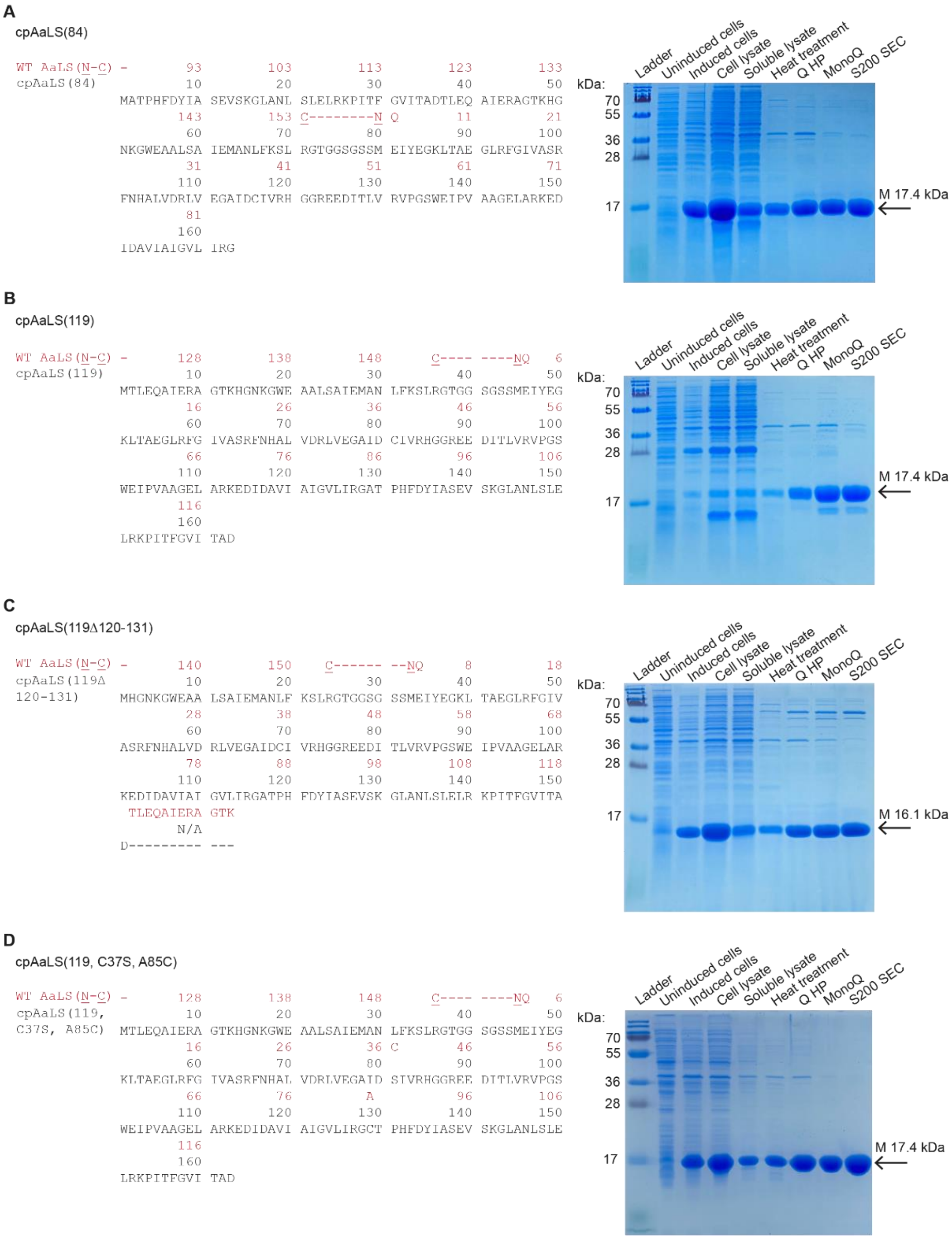


**Fig. S16. Primary sequence and purification of cpAaLS variants.**

(A-D) Primary sequence (left) and SDS-PAGE analysis (right) of cpAaLS(84) (A), cpAaLS(119) (B), cpAaLS(119Δ120-131) (C), and cpAaLS(119, C37S, A85C) (D) variants. Each primary sequence (in black) is aligned to wild-type AaLS (in red) showing its native termini (N-C), gaps (-), amino acid substitutions (i.e. C37C), or truncations (Δ120-131). The arrows next to the respective gels indicate the band corresponding to the molecular mass (M in kDa) calculated from the amino acid sequence using the Expasy ProtParam tool (*58*).

**Table S1. Oligonucleotides used in this study.**

| Name | Sequence |
|---|---|
| FW_NdeI_cpAaLS(119Δ120-131) | CATCATATGCACGGCAACAAAGGTTGG |
| RV_XhoI_cpAaLS(119Δ120-131) | CATCATCTCGAGTTAGTCAGCTGTAATAAC |
| FW_cpAaLS(119, C37S) | CGTCTGGTGGAGGGTGCAATTGATaGCATAGTCCGTCATGGCGGC |
| RV_cpAaLS(119, C37S) | GCCGCCATGACGGACTATGCtATCAATTGCACCCTCCACCAGACG |
| FW_cpAaLS(119, C37S, A85C) | CAATTGGCGTTCTCATCAGAGGCTGCACGCCACATTTCGATTATATCGCC |
| RV_cpAaLS(119, C37S, A85C) | GGCGATATAATCGAAATGTGGCGTGCAGCCTCTGATGAGAACGCCAATTG |

**Table S2. Plasmids used in this study.**

| Name | Gene | Promoter/Operator[a] | Ori | Marker[b] | Ref. |
|---|---|---|---|---|---|
| pMG_cpAaLS_L8 (119) | cpAaLS (119) | $P_{T7}$ / *lac*O | pBR322 | $Amp^R$ | 39 |
| pMG_cpAaLS_L8 (84) | cpAaLS (84) | $P_{T7}$ / *lac*O | pBR322 | $Amp^R$ | 41 |
| pMG_cpAaLS_L8 (119Δ120-131) | cpAaLS (119Δ120-131) | $P_{T7}$ / *lac*O | pBR322 | $Amp^R$ | This study |
| pMG_cpAaLS_L8 (119, C37S) | cpAaLS (119, C37S) | $P_{T7}$ / *lac*O | pBR322 | $Amp^R$ | This study |
| pMG_cpAaLS_L8 (119, C37S, A85C) | cpAaLS (119, C37S, A85C) | $P_{T7}$ / *lac*O | pBR322 | $Amp^R$ | This study |

[a] $P_{T7}$ / *lac*O, T7 promoter combined with lactose operator.

[b] $Amp^R$, ampicillin resistance.

### Table S3. Cryo-EM data collection and structure refinement statistics.

| **Variant** | cpAaLS(84) | cpAaLS(119) | cpAaLS(119) | cpAaLS(119) | cpAaLS(119, C37S, A85C) | | |
|---|---|---|---|---|---|---|---|
| **Assembly** | 12-pentamer spherical cage | 24-pentamer spherical cage | 36-pentamer spherical cage | Straight tube | Twisted tube 28-Å gap | Twisted tube 18-Å gap | Twisted tube 0-Å gap |
| **EMDB** | 51006 | 51005 | 51004 | 51003 | 51001 | 51000 | 50999 |
| **PDB** | 9G3P | 9G3O | 9G3N | 9G3M | 9G3J | 9G3I | 9G3H |
| **Data collection and processing** | | | | | | | |
| Magnification | 105,000 × | 105,000 × | 105,000 × | 81,000 × | 105,000 × | 105,000 × | 105,000 × |
| Voltage (keV) | 300 | 300 | 300 | 300 | 300 | 300 | 300 |
| Electron exposure ($e^-/Å^2$) | 40 | 40 | 40 | 40 | 42.44 | 42.44 | 42.44 |
| Defocus range | -0.9 to -2.1 | -0.9 to -2.1 | -0.9 to -2.1 | -0.9 to -3.0 | -0.9 to -1.5 | -0.9 to -1.5 | -0.9 to -1.5 |
| Pixel size (Å) | 0.86 | 0.86 | 0.86 | 1.1[a] | 0.846 | 0.846 | 0.846 |
| Micrographs (no.) | 8,486 | 10,060 | 9,241 | 5,031 | 12,746 | 12,746 | 12,746 |
| Symmetry imposed | I | T | T | 81.6 º twist 64.4 Å rise | 57.0 º twist 24.5 Å rise | 57.3 º twist 22.1 Å rise | 57.7 º twist 19.8 Å rise |
| Initial particle images (no.) | 590,975 | 374,850 | 46,386 | 171,841 | 1,620,295 | 1,620,295 | 1,620,295 |
| Final particle images (no.) | 98,492 | 83,496 | 22,183 | 168,323 | 192,310 | 807,640 | 208,379 |
| Map resolution (Å) | 2.08 | 2.76 | 3.07 | 3.09 | 3.10 | 2.85 | 2.44 |
| FSC threshold | 0.143 | 0.143 | 0.143 | 0.143 | 0.143 | 0.143 | 0.143 |
| Map resolution range (Å) $25^{th}$–$75^{th}$ percentile | 2.0–2.3 | 2.7–3.2 | 3.0–4.0 | 3.0–4.0 | 2.9–4.5 | 2.6–3.7 | 2.3–3.2 |
| Map sharpening B-Factor | 65.0 | 99.3 | 87.9 | 107.1 | 102.5 | 115.2 | 79.6 |
| **Model building** | | | | | | | |
| PDB code of the initial model: 1HQK | | | | | | | |
| Model composition | | | | | | | |
| Chains | 60 | 120 | 180 | 150 | 100 | 100 | 100 |
| Protein residues | 9600 | 17856 | 26652 | 23010 | 15260 | 15260 | 15260 |
| Non-hydrogen atoms | 72480 | 133824 | 199560 | 172740 | 114640 | 114640 | 114640 |
| Water/Ligands | 0 | 0 | 0 | 0 | 0 | 0 | 0 |
| Nucleotides | 0 | 0 | 0 | 0 | 0 | 0 | 0 |
| *B* factors (mean $Å^2$) | 32.62 | 50.76 | 96.35 | 74.82 | 118.52 | 110.28 | 78.30 |
| R.M.S. deviations | | | | | | | |
| Bond lengths (Å) | 0.004 | 0.005 | 0.004 | 0.004 | 0.004 | 0.004 | 0.005 |
| Bond angles (º) | 0.963 | 0.974 | 0.943 | 0.945 | 0.926 | 0.915 | 0.994 |
| Validation | | | | | | | |
| MolProbity score | 0.98 | 1.17 | 1.22 | 1.07 | 1.22 | 1.18 | 1.21 |
| Clash score | 2.10 | 1.69 | 2.62 | 1.87 | 2.56 | 3.60 | 2.79 |
| Poor rotamers (%) | 0.11 | 0.00 | 0.06 | 0.00 | 0.00 | 0.00 | 0.00 |
| Ramachandran | | | | | | | |
| Favored (%) | 98.10 | 96.28 | 97.04 | 97.38 | 96.95 | 97.88 | 97.21 |
| Allowed (%) | 1.90 | 3.72 | 2.96 | 2.62 | 3.05 | 2.12 | 2.79 |
| Outliers (%) | 0.00 | 0.00 | 0.00 | 0.00 | 0.00 | 0.00 | 0.00 |
| CC (volume) | 0.86 | 0.78 | 0.77 | 0.76 | 0.86 | 0.86 | 0.90 |

[a] Super-resolution pixel size 0.55 Å
